# Dysregulation of an H3K79me2-dependent epigenetic barrier impairs neural progenitor cell proliferation and differentiation in Fragile X syndrome

**DOI:** 10.1101/2025.08.31.673358

**Authors:** Olivier Dionne, Gabrielle-Lee Gaudreau, Clara Bergis-Ser, Mariano Avino, Salomé Sabatié, François Corbin, Benoit Laurent

**Affiliations:** Department of Biochemistry and Functional Genomics, Faculty of Medicine and Health Sciences, Université de Sherbrooke, Sherbrooke, QC, Canada; Research Center on Aging, Centre Intégré Universitaire de Santé et Services Sociaux de l’Estrie-Centre Hospitalier Universitaire de Sherbrooke, Sherbrooke, QC, Canada; Bio-informatic platform, Department of Biochemistry and Functional Genomics, Faculty of Medicine and Health Sciences, Université de Sherbrooke, Sherbrooke, QC, Canada; Centre de Recherche du Centre Hospitalier Universitaire de Sherbrooke (CRCHUS), Sherbrooke, QC, Canada

**Keywords:** Fragile X syndrome, FMRP, epigenetic, neurodevelopment, differentiation, neural progenitor cells

## Abstract

Fragile X syndrome (FXS) is a neurodevelopmental disorder caused by silencing of the *FMR1* gene, which encodes the multifunctional RNA-binding protein FMRP. While FMRP is best known for its roles in RNA metabolism, it can also associate with chromatin through recognition of histone H3 lysine 79 di-methylation (H3K79me2), an epigenetic mark linked to transcriptionally active genes. However, the functional relevance of this FMRP–H3K79me2 interaction has remained largely unexplored in the context of FXS pathophysiology. We assessed H3K79me2 levels during the differentiation of induced pluripotent stem cells (iPSCs) generated from both healthy individuals and FXS patients and discovered a global increase in H3K79me2 levels specifically in FXS neural progenitor cells (NPCs). Altered H3K79me2 landscape drives widespread transcriptional dysregulation, characterized by reduced expression of neurogenesis-associated genes alongside aberrant activation of programs promoting proliferation and glial lineage commitment. Functionally, these changes result in enhanced NPCs proliferation and a biased differentiation trajectory. Notably, pharmacological reduction of global H3K79me2 levels in FXS NPCs using DOT1L inhibitors effectively mitigates these defects, restoring proliferation rate and rebalancing lineage specification, thereby rescuing key aspects of the pathological phenotype. Collectively, our findings identify an H3K79me2-dependent epigenetic barrier regulating NPC proliferation and lineage commitment and link its dysregulation to the neurodevelopmental defects associated with FXS.

## Introduction

Fragile X syndrome (FXS) is a neurodevelopmental disorder representing the leading inherited cause of intellectual disability, as well as the most prevalent monogenic contributor of autism spectrum disorders (ASD)^1–3^. FXS results from an expansion of CGG trinucleotides repeats found within the 5’ untranslated region (UTR) of the Fragile X Messenger Ribonucleoprotein 1 (FMR1) gene. Expansion beyond the pathological threshold of 200 repeats leads to methylation of nearby CpG islands and subsequent transcriptional silencing of FMR1^4–6^. Thus, the pathophysiological mechanisms of FXS are fundamentally driven by the loss of expression of the protein encoded by FMR1, the Fragile X messenger ribonucleoprotein (FMRP), formerly known as the Fragile X mental retardation protein^7^.

FMRP is a widely expressed RNA-binding protein predominantly found in the brain and testis^8–10^. It is composed of at least three RNA-binding domains: two distinct nuclear ribonucleoprotein K homology domains (KH1 and KH2) and one arginine-glycine-glycine (RGG) box located within the C-terminus^11^. Given its RNA-binding capability, FMRP is involved in a large range of RNA-based molecular processes, including RNA editing, splicing, stability and translation^12,13^. However, regulation of target mRNA translation is widely recognized as the canonical function of FMRP^14,15^. Thus, FMRP absence has been repeatedly linked to a global disruption of protein synthesis rates across various human cellular models^16–18^.

FMRP also contains a nuclear localisation signal (NLS), a nuclear export signal (NES) as well as two N-terminally located Agenet domains that mediate FMRP interactions with chromatin and other proteins^19,20^,. FMRP can directly interact via its Agenet domains with histone H3 lysine 79 di-methylation (H3K79me2), a histone mark enriched at transcriptionally active genes^21,22^. Interestingly, FMRP point mutations in NLS or NES have been associated with clinical features resembling those of FXS, including intellectual disability, developmental delay, and seizures^23–26^. Although FMRP is predominantly localized in the cytoplasm, the loss of its nuclear functions may play a significant role in FXS pathophysiology^27–29^.

*Fmr1* knock-out (KO) animal models have been extensively used to investigate the pathophysiology of FXS and to develop targeted pharmacological interventions^30,31^. However, the consistent failure of these candidate drugs in clinical trials highlights the limited translational relevance of animal-based findings and suggests that these models do not fully replicate FXS human phenotypes^32–35^.

Brain development depends on the coordination of tightly regulated processes, including cell proliferation, specification, and maturation^36^. While the overall sequence of these events is conserved across mammals, human neurodevelopment proceeds at a markedly slower pace than in rodents, contributing to the distinctive complexity and architecture of the human brain^37–39^. This protracted developmental timeline is maintained *in vitro*, as observed during the differentiation of human induced pluripotent stem cells (iPSCs)^40–42^. The transition from iPSCs to neural progenitor cells (NPCs) and ultimately mature neural cells, such as neurons and glial cells, is driven by intrinsic molecular programs that orchestrate the sequential progression of these developmental processes.

Recent studies have revealed that epigenetic mechanisms are critical for regulating the orderly progression of neural cells through a series of distinct developmental stages on a highly precise timeline^43,44^. In progenitor cells, an epigenetic barrier holds transcriptional programs in a poised state, which must be lifted for neurons to acquire mature properties upon differentiation^45^. Furthermore, a recent study demonstrated that RNA-binding proteins mediate NPCs differentiation trough interaction with the chromatin^46^. Given FMRP ability to bind chromatin, we hypothesized that it participates in the epigenetic regulation of neural differentiation, a process potentially disrupted in FXS. Using an iPSCs-based model derived from individuals with FXS, we identified neurodevelopmental defects in NPCs and showed that FMRP loss impairs critical progenitor-stage molecular events, thereby hindering their differentiation trajectory.

## Results

### Stage-specific dysregulation of H3K79me2 global levels in FXS NPCs

FMRP has been reported to bind chromatin via direct recognition of H3K79me2^22^, leading us to hypothesize that it may contribute to the epigenetic regulation of neural differentiation. To test this, we examined global H3K79me2 levels at sequential stages of human iPSCs differentiation into NPCs and neural cultures, following established protocols (**Figure 1A**)^47,48^. Our study included four control (CTL) and four FXS iPSCs lines including one FMR1-KO line and three iPSCs lines derived from FXS patients that we have previously generated and validated^49–51^ (**Supplementary Table 1**). Global H3K79me2 levels were comparable between CTL and FXS iPSCs while FXS cells exhibited around 1.75-fold increase of H3K79me2 levels at the NPC stage, relative to controls (**Figure 1B**). At day 30 of a neural culture, this difference was no longer detectable (**Figure 1B**). Stage-to-stage analysis revealed that CTL lines showed a marked increase in H3K79me2 (6-fold increase at NPCs stage, 14-fold increase in neural cultures) (**Supplementary Figure 1A**), consistent with previous observations during neuronal differentiation of mouse embryonic stem cells^52^. In contrast, FXS lines showed a 14-fold increase in H3K79me2 levels during the iPSC-to-NPC transition, with levels remaining stable thereafter in neural cultures (**Supplementary Figure 1B**). These results indicate a developmental stage–specific disruption of global H3K79me2 regulation in the absence of FMRP, confined to the NPCs stage of differentiation.

**Figure 1.**
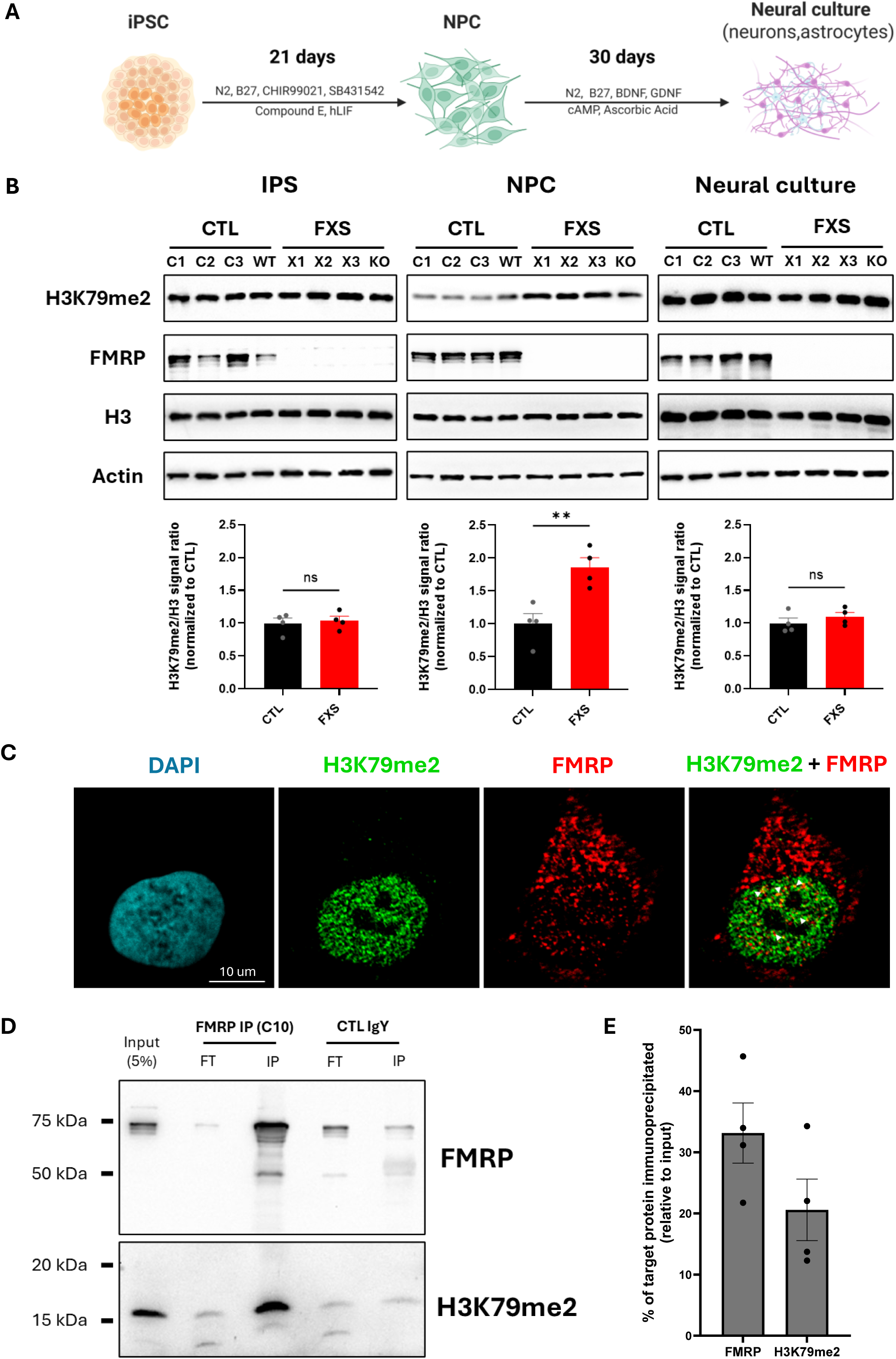
Developmental stage-specific dysregulation of H3K79me2 in FXS NPCs. **(A)** Schematic of the differentiation protocols used to generated neural progenitor cells (NPCs) and neural mature culture from induced pluripotent stem cells (iPSCs). **(B)** Protein levels of H3K79me2 across the different stages of the differentiation model were analyzed by western-blot (top panel). On the bottom panel, quantification of H3K79me2 global levels (n =4/genotype). Results are indicated as mean ± SEM, **p < 0.01. **(C)** Representative immunofluorescence image showing FMRP nuclear localization and co-localization with H3K79me2. **(D)** Representative immunoblots of FMRP co-immunoprecipitation with H3K79me2 within Benzonase nuclease-treated nuclear fractions of NPCs. This representative experiment was performed in the NPC_C3 line. The other replicates can be found in **Supplementary Figure 1. (E)** Quantification of the proportion of target proteins enrichment following FMRP immunoprecipitation in the nuclear fraction of NPCs (n= 4 cell lines). Results are expressed as a percentage relative of the input sample. Results are indicated as mean ± SEM. ** p<0.01,

Since FMRP can directly bind this histone modification in vitro^22^, we first examined by immunofluorescence whether FMRP colocalizes with H3K79me2 in the nuclei of NPCs. At this stage of differentiation, FMRP immunostaining confirmed both cytoplasmic and nuclear localization (**Figure 1C**). Merged FMRP and H3K79me2 signals suggested multiple nuclear regions of colocalization within NPCs nuclei. To validate this observation, we assessed FMRP capability to bind chromatin, specifically H3K79me2 modification, by performing FMRP immunoprecipitation from NPCs nuclear-enriched fractions. This assay confirmed the interaction between FMRP and H3K79me2 (**Figure 1D**; **Supplementary Figure 1C**) and showed that approximately 20% of all H3K79me2 can be immunoprecipitated by FMRP (**Figure 1E**). Collectively, these findings demonstrate that FMRP interacts with H3K79me2 in NPCs and suggest that the loss of FMRP - and thus the loss of this interaction - may specifically disrupt the regulation of this epigenetic mark in FXS NPCs.

### Alterations in the H3K79me2 landscape drive gene expression changes in FXS NPCs

H3K79me2 is strongly enriched in transcriptionally active genes, with high levels located near the transcription starting site (TSS) that gradually decline throughout the gene body^21^. However, its level often correlates with transcriptional elongation rather than initiation^53,54^. Hence, H3K79me2 misregulation in FXS NPCs could lead to aberrant gene activation in these cells. To address this point, we examined the H3K79me2 landscape by ChIP-seq experiments against this histone mark in both control and NPCs and assessed in parallel global gene expression by RNA-seq in the same cells. First, the principal component analysis (PCA) of ChIP-seq data showed that the global H3K79me2 landscape in control NPCs was very similar between our four control NPC populations (**Figure 2A**). H3K79me2 landscape in FXS NPCs also had high consistency between the four populations but was distinct from that of control NPCs as indicated by the average position of the samples on the x-axis which represents the first principal component responsible for 60.9% of the variance (**Figure 2A**). ChIP-seq analysis further identified a total of 7,339 peaks with a differential H3K79me2 abundance in FXS NPCs compared to control NPCs. Among these differentially enriched peaks, 4,190 regions were identified with an increase in H3K79me2 abundance and 3,149 with a decreased enrichment (**Figure 2B**). These results confirm the disruption of global H3K79me2 landscape in FXS NPCs.

**Figure 2.**
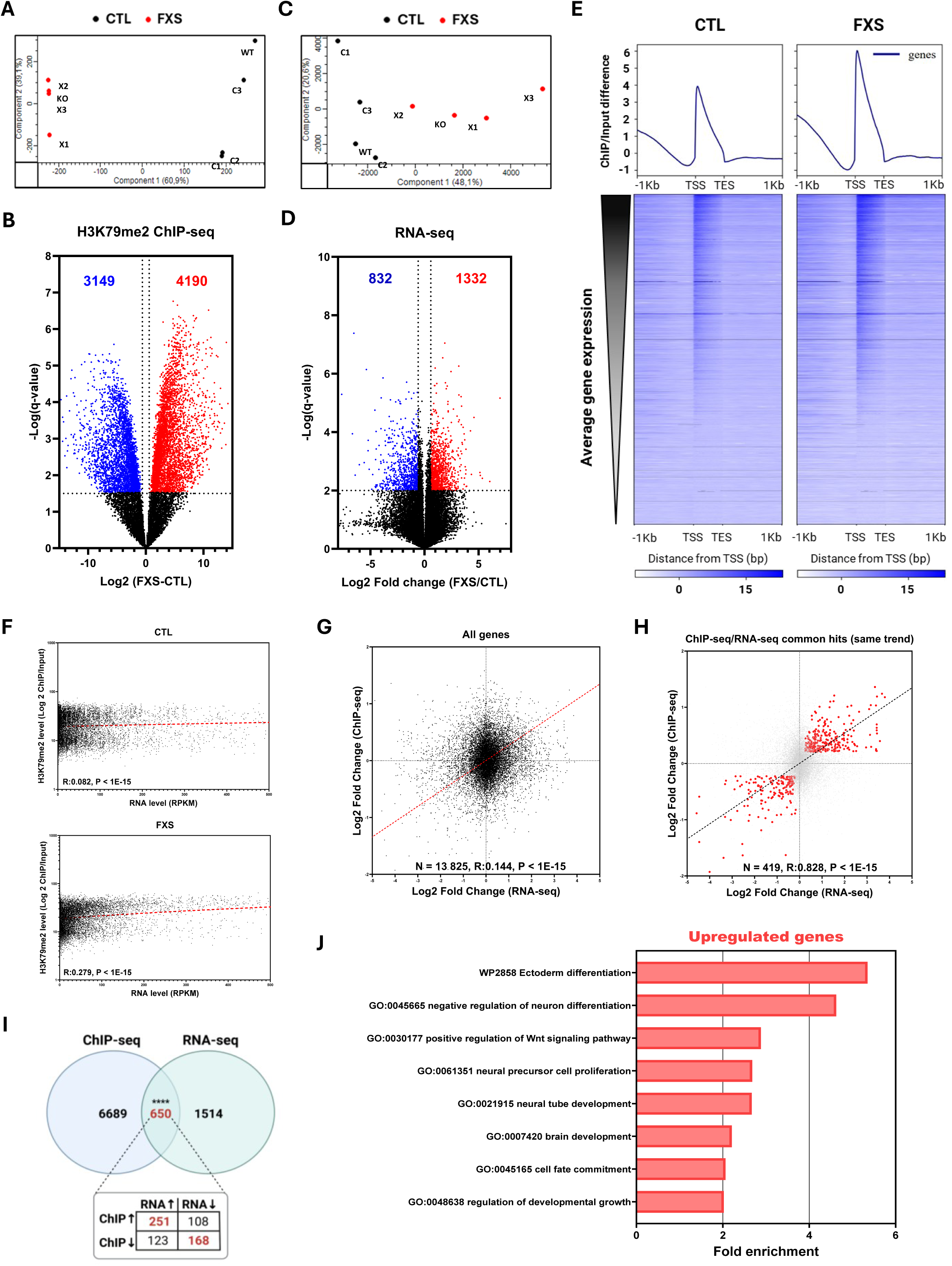
Aberrant H3K79me2 deposition and gene expression patterns in FXS NPCs. **(A)** Principal component analysis showing clustering of the samples based on genotype for the ChIP-seq experiments. **(B)** Volcano plot of the -log10 (q-value) versus the log2 of the difference between H3K79me2 level (reported as log2 of ChIP/input signal ratio) in control (CTL) and FXS NPCs. Peaks showing decreased abundance in FXS are labeled in blue, and peaks showing increase abundance are labeled in red. **(C)** Principal component analysis showing clustering of the samples based on genotype for the RNA-seq experiments. **(D)** Volcano plot of -log10 (q-value) versus the log2 of Fold change for the RNA-seq dataset. Genes with increased expression (Log2 Fold change ˃ 0.58) are labeled in red, and genes with reduced expression (Log2 Fold change < -0.58) in blue. **(E)** Metaplots of average H3K79me2 signal (expressed as ChIP/input difference) along gene bodies in CTL and FXS NPCs (top panel) and heatmaps of H3K79me2 enrichment along gene bodies, ranked by gene expression levels determined by RNA-seq (n= 13,825 genes) (bottom panel). **(F)** Correlation analysis of H3K79me2 abundance (determined by the ChIP/input signal ratio) and RNA expression quantified by RNA-seq for corresponding genes in CTL and FXS NPCs. **(G-H)** Correlation analysis between fold change of H3K79me2 abundance and fold change of RNA levels in FXS NPCs in corresponding genes (n= 13,825) for **(G)** all genes commonly identified between the ChIP-seq and RNA-seq datasets and **(H)** for genes found dysregulated in both ChIP and RNA-seq datasets with the same trend of perturbation. **(I)** Venn diagram showing the number H3K79me2 peaks with differential abundance, the number of differentially expressed genes and the overlap between the two datasets in FXS NPCs. Fisher’s exact test, **** p<0.00001. **(J)** (A) Representation of functional annotations enriched among upregulated genes identified in FXS NPCs.

For the RNA-seq, the PCA analysis showed a similar genotype-based clustering, with more variability for the four FXS NPCs populations with a 41.8% average variance (**Figure 2C**). Data analysis revealed a total of 2,164 differentially expressed genes (DEGs) in FXS NPCs compared to CTL NPCs. Among these DEGs, 1,332 genes (61.5%) were upregulated in FXS NPCs while 832 genes (38.5%) were downregulated (**Figure 2D**). As H3K79me2 levels correlate with gene expression, we next performed a metaplot analysis of H3K79me2 signal profile along the gene body and surrounding regions (± 1 kb) for each gene and ranked these profiles based on the level of gene expression detected by RNA-seq (total of 13,825 genes). The genomic distribution of H3K79me2 in both genotypes followed the expected pattern, with an enrichment peaking around the TSS and gradually decreasing along the gene body (**Figure 2E**, top panel). Higher H3K79me2 levels were detected in FXS NPCs compared to CTL NPCs, consistent with the global increase of this epigenetic mark observed by western blot (**Figure 1B**). Ranking of H3K79me2 signal profiles based on the gene expression showed higher enrichment in this histone mark for the most highly expressed genes in both genotypes, consistent with previous reports^21,55^ (**Figure 2E**).

We next examined how H3K79me2 enrichment related to transcript abundance within genes. By plotting H3K79me2 levels against RNA expression and performing correlation analysis, we found that, in control NPCs, the association was weak (r = 0.082; **Figure 2F**, left panel). In contrast, FXS NPCs showed a 3-fold higher correlation (r = 0.279), indicating that gene expression in FXS cells was more closely coupled to the H3K79me2 landscape (**Figure 2F**, right panel). These observations prompted us to hypothesize that the transcriptional changes in FXS NPCs may, at least in part, result from altered H3K79me2 levels. To test this, we compared fold changes in H3K79me2 enrichment with corresponding fold changes in gene expression across all genes present in both the ChIP-seq and RNA-seq datasets. At the global level, this analysis only revealed a weak correlation (r = 0.144) (**Figure 2G**). However, when the analysis was focused on genes consistently dysregulated in both datasets with concordant changes in H3K79me2 and RNA expression, we observed a remarkably strong correlation (r = 0.828) (**Figure 2H**). Integration of ChIP-seq (H3K79me2 enrichment changes) and RNA-seq (gene expression changes) datasets revealed a significant overlap of 650 commonly dysregulated genes. (**Figure 2I**). Among these genes, 251 had both increase in H3K79me2 level and gene expression while 168 genes had both decrease in H3K79me2 level and gene expression. These two groups (n= 419 genes) represented cases where H3K79me2 changes were concordant with gene expression changes, consistent with a positive correlation. The remaining genes (n= 231 genes) showed discordant patterns (e.g., increased H3K79me2 but decreased expression, or vice versa) (**Figure 2I**). The majority of these genes (419/650 genes = 64.4%) showed directionally consistent changes, suggesting that alterations in H3K79me2 enrichment were closely associated with transcriptional regulation in FXS NPCs. These findings support the idea that dysregulated H3K79me2 contributes to the gene expression changes observed.

### Dysregulated H3K79me2-associated genes control core NPC functional pathways

We next assessed the functional relevance of the genes whose alterations in H3K79me2 enrichment strongly correlated with changes in transcript expression in FXS NPCs. A Gene Ontology (GO) enrichment analysis revealed that upregulated genes with higher H3K79me2 levels were enriched for biological processes that are essential for NPC proliferation and critical for establishing NPC identity (**Figure 2J**). From among the top enriched genes, we selected four candidates known to regulate NPCs proliferation and differentiation (PAX6), or involved in key signaling cascades such as the Wnt (WNT7A), Notch (DLL1), and fibroblast growth factor (FGFR2) pathways^56–63^. We confirmed that all four genes displayed increased H3K79me2 enrichment along with elevated mRNA expression in FXS NPCs relative to controls (**Supplementary Figure 2A-B**). Notably, upregulation of PAX6 has been previously shown to enhance NPCs proliferation at the expense of differentiation during corticogenesis^60,64^, consistent with the enrichment of the GO term *Negative regulation of neuron differentiation* (**Figure 2J**). We next examined additional candidate genes involved in neurogenesis (SLIT1, PIEZO1)^65–67^ and gliogenesis (SOX8, SOX9)^68–70^. In FXS NPCs, neurogenesis-related genes exhibited reduced H3K79me2 levels and lower transcript abundance, whereas critical regulators of gliogenesis showed increased H3K79me2 enrichment and elevated RNA expression (**Supplementary Figure 2C-F**). Together, these findings suggest that FXS NPCs may display enhanced proliferation, impaired differentiation potential, and a bias toward aberrant cell fate acquisition during differentiation.

### Increased proliferation of FXS NPCs

Transcriptomic data indicated upregulation of genes involved in cell proliferation in FXS NPCs. We first assessed the proliferation of CTL and FXS NPCs and found that FXS progenitors proliferated faster than control ones (**Figure 3A**). This higher proliferative capacity was further confirmed by colony-forming assays (**Figure 3B**). We next performed immunofluorescence staining for the proliferation marker Ki67 and showed a higher proportion of proliferating cells in the FXS population than in the CTL one, as indicated by the percentage of Ki67-positive cells (37.60±3.50 % vs 16.85±1.48 % normalized to DAPI^+^ cells or 39.86±3.71 % vs 17.69±1.55 % normalized to SOX2^+^ cells) (**Figure 3C**). To gain insight into the mechanisms driving the increased proliferation of FXS NPCs, we investigated the regulation of cell cycle progression. Analysis of cell cycle phase distribution using propidium iodide (PI) staining followed by fluorescence-activated cell sorting (FACS) revealed that the FXS NPCs population had a higher proportion of cells in the G2/M phases compared to the CTL NPCs population (9.57±1.21 % vs 4.42±0.38 %) (**Figure 3D**). Moreover, analysis of cell cycle kinetics following synchronization at the G2/M boundary using nocodazole^71^ showed that FXS NPCs progressed more rapidly through this phase (**Figure 3E**). Approximately 50 % of cells synchronized with nocodazole exited G2/M cell cycle phase 2h after nocodazole removal (**Figure 3F**). Consistently, immunofluorescence staining for phosphorylated histone H3 at serine 10 (pH3), a marker of mitosis, showed a higher proportion of FXS NPCs in M phase compared to controls (**Supplementary Figure 3A**).

**Figure 3.**
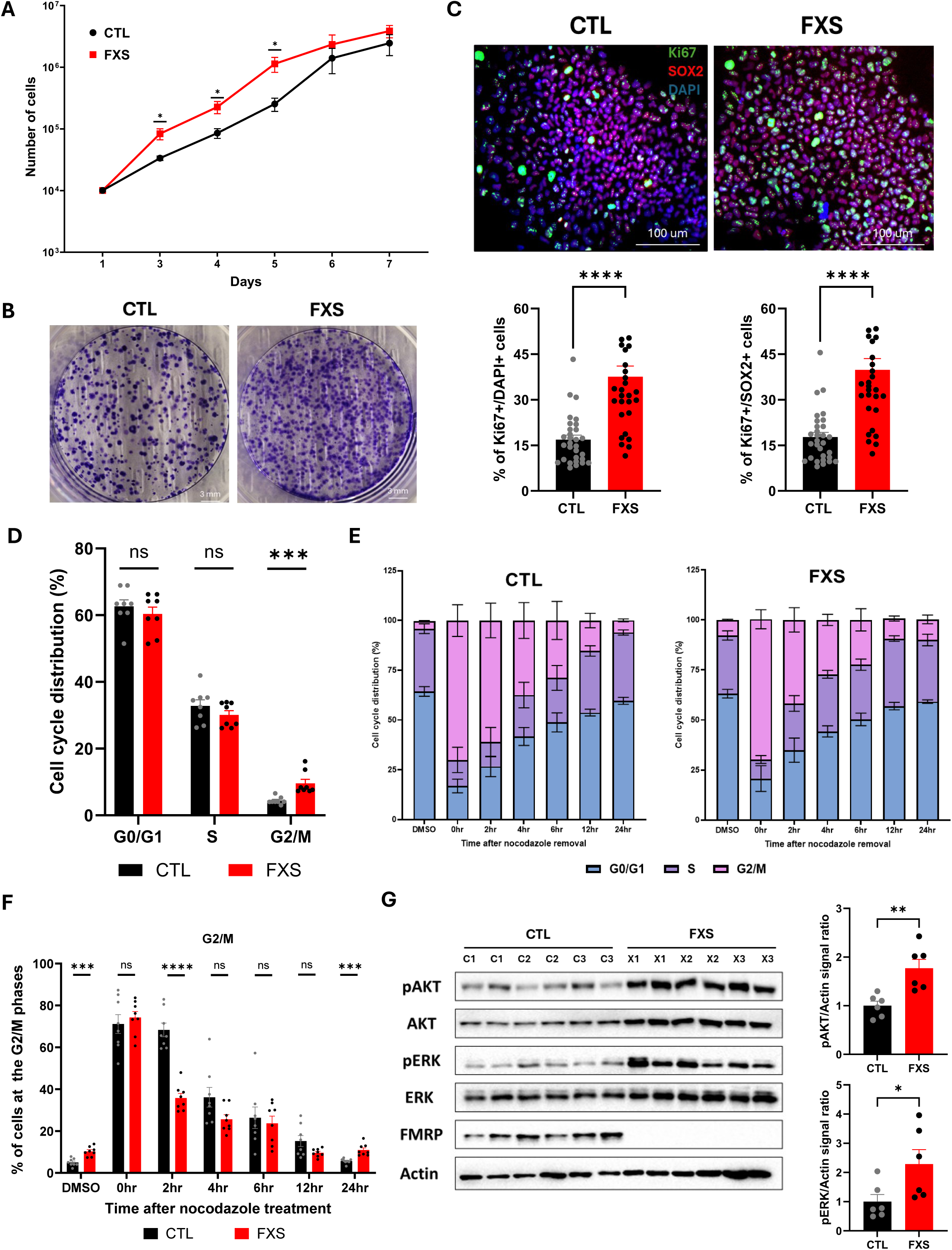
Increased proliferation and impaired cell cycle in FXS NPCs. **(A)** Growth curves of control (CTL) and FXS NPCs over 7 days (n= 4 cell lines/genotype; 2 replicates/cell line). **(B)** Representative colony-forming assays of CTL and FXS NPCs. **(C)** Representative immunostaining for the proliferation marker Ki67 in CTL and FXS NPCs (top panel), with quantification of Ki67^+^ cells normalized to either DAPI^+^ or SOX2^+^ cell counts (n= 4 cell lines/genotype, 5 fields/cell line) (bottom panel). **(D-E)** Cell cycle phase distribution analyzed by Fluorescence-Activated Cell Sorting (FACS) after DNA staining with propidium iodide in **(D)** unsynchronized NPCs and **(E)** NPCs released from nocodazole-induced synchronization (n = 4 cell lines/genotype; 2 replicates per cell line). **(F)** Quantification of the proportion of NPC at the G2/M phase following release of the nocodazole-induced synchronization (n = 4 cell lines/ genotype; 2 replicates per cell line). **(G)** Protein levels of ERK and AKT activation following synchronization with the CDK1 inhibitor RO-3306 were analyzed by western-blot (left panel). On the right panel, quantification of phospho-ERK and phospho-AKT levels (n= 3 cell lines/genotype; 2 replicates/cell line). Results are indicated as mean ± SEM. *p< 0.05, ** p<0.01, ***p<0.0001, ****p<0.00001.

Major signaling kinases such as protein kinase B (AKT) and extracellular signal-regulated kinase (ERK), integrate extracellular growth signals and regulate progression through different phases of the cell cycle. Both AKT and ERK have been consistently reported to be hyperactivated in FXS^18,72–74^. We next examined the activation status of these kinases in FXS NPCs at the G2/M boundary. To do so, we synchronized control and FXS NPCs at the G2/M boundary using nocodazole or the CDK1 inhibitor RO-3306^71,75^. As a control, NPCs were also synchronized at the G1 phase using the CDK4/6 inhibitor PD 0332991^76^ (**Supplementary Figure 3B**). We showed an increased phosphorylation of both ERK and AKT in FXS NPCs synchronized in G2/M phase either with RO-3306 inhibitor (**Figure 3G**) or with nocodazole (**Supplementary Figure 3C**). G1 synchronization of NPCs did not reveal any significant differences for AKT and ERK activation (**Supplementary Figure 3D**). Together, these findings indicate that the higher proliferation of FXS NPCs may result from dysregulated G2/M phase transition driven by increased activation of the ERK and AKT signaling pathways.

### FXS NPCs exhibit abnormal differentiation trajectory

In addition to increased proliferation, transcriptomic data indicated that FXS NPCs may have compromised differentiation potential, predisposing them to abnormal cell fate. To verify this, we first evaluated NPCs efficiency to differentiate into a mature neural culture over a 30-day differentiation period. At day 30 (D30), immunofluorescence staining for the neuronal cytoskeletal markers MAP2 and TUBB3 showed that FXS NPCs exhibited impaired neuronal differentiation, as evidenced by the reduced MAP2⁺ (46.09 ± 6.02 mm^2^ vs 109.30 ± 7.37 mm^2^) and TUBB3⁺ (54.14± 9.36 mm^2^ vs 121.4± 6.71 mm^2^) masked area normalized by the number of DAPI+ cells relative to CTL (**Figure 4A**). We next quantified transcript levels of neuronal lineage-specific markers across the differentiation period. Expression levels of neuronal markers NeuN, MAP2, and GAP43 showed a robust and early upregulation in CTL lines, whereas FXS lines exhibited a markedly reduced induction of these markers (**Figure 4B; Supplementary Figure 4A**), suggesting a compromised capacity to acquire mature neuronal identity. We also analyzed PLZF, an NPC-specific marker, and showed a rapid and marked decreased expression during the differentiation of CTL NPCs, confirming their commitment toward differentiation (**Figure 4B**). In contrast, PLZF expression in FXS NPCs declined more gradually and to a lesser extent, confirming impaired differentiation potential (**Figure 4B**).

**Figure 4.**
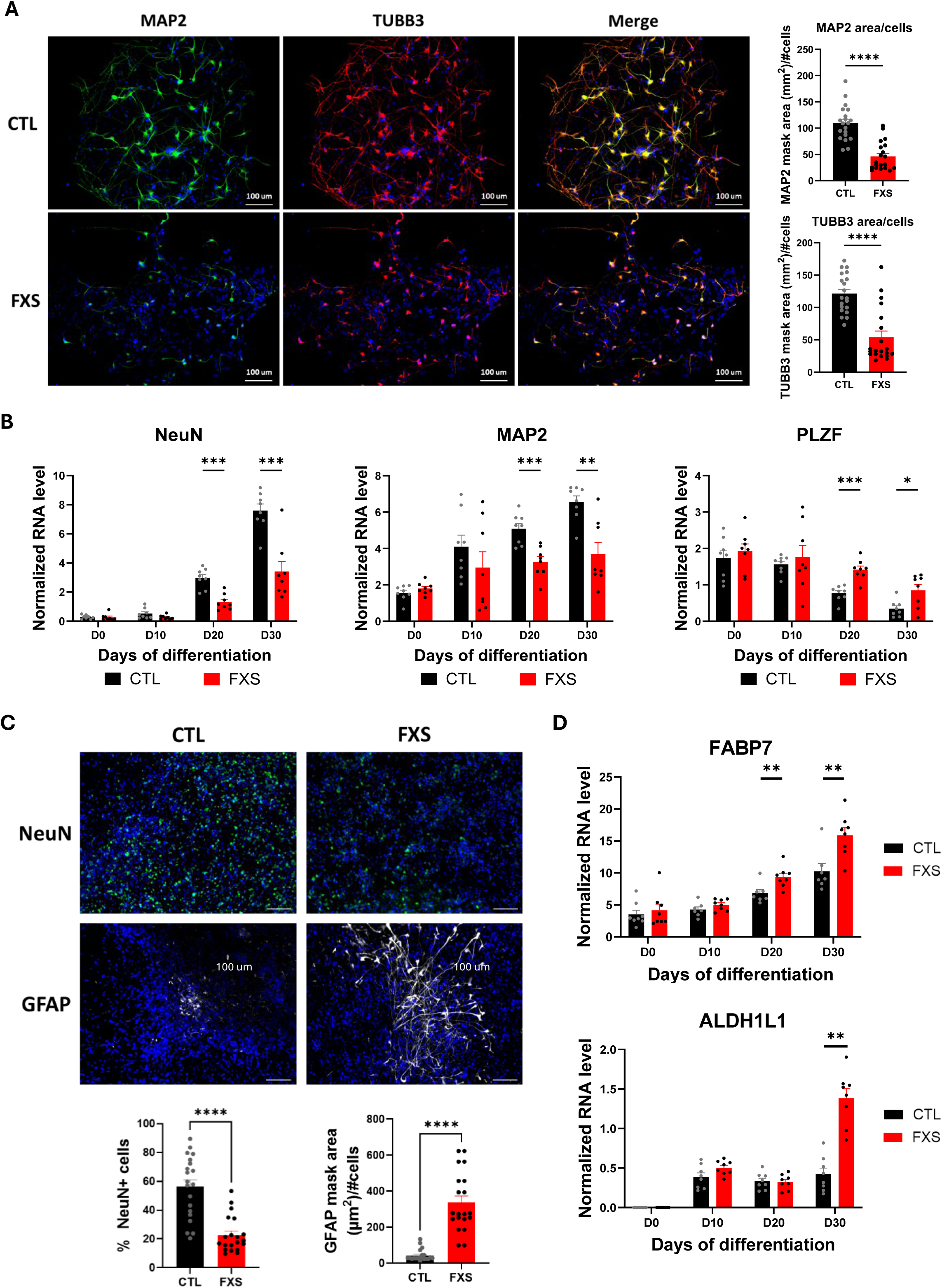
Aberrant differentiation trajectory in FXS NPCs. **(A)** Representative immunostaining against the neuronal cytoskeleton markers MAP2 and TUBB3 in control (CTL) and FXS in neural culture at day 30 (D30) (left panel). On the right panel, quantification of the MAP2 and TUBB3 masked area normalized by the DAPI^+^ cell count (n= 4 cell lines/genotype, 5 fields/cell line) **(B)** Quantification of neuronal lineage-specific and NPC-specific markers by quantitative PCR during the differentiation of CTL and FXS NPCs into neural culture (n= 4 cell lines/genotype, 2 replicates/cell line). **(C)** Representative immunostaining against the neuron marker NeuN (quantified by the proportion of NeuN^+^ cells normalized by the DAPI^+^ cell count) and the astrocyte marker GFAP (quantified by the GFAP masked area normalized by the DAPI^+^ cell count) in a D30 neural culture (n = 4 cell lines/genotype, 5 fields/cell line). **(D)** Quantification of astrocyte-specific markers by quantitative PCR during the differentiation of CTL and FXS NPCs into neural culture (n = 4 cell lines/genotype, 2 replicates/cell line). Results are indicated as mean ± SEM. *p< 0.05, ** p<0.01, ***p<0.0001, ****p<0.00001.

Transcriptomic profiling revealed that in FXS NPCs, neurogenesis-related genes (SLIT1, PIEZO1) were downregulated, whereas gliogenesis-associated genes (SOX8, SOX9) were upregulated (**Supplementary Figure 2C-F**), suggesting a bias toward aberrant cell fate during differentiation. To test this, we assessed NPCs differentiation into astrocytes by immunofluorescence staining for the astrocytic marker GFAP, with NeuN as a neuronal control. By day 30 (D30), FXS cultures displayed a marked increase in GFAP⁺ masked aera normalized by the number of cells compared to CTL (337.8 ± 36.55 mm^2^ vs. 43.26 ± 7.29 mm^2^) (**Figure 4C**), indicating a heightened propensity for astrocytic differentiation. Consistently, NeuN staining confirmed impaired neuronal differentiation within the same cultures (**Figure 4C**). Elevated expression of astrocytic markers FABP7, ALDH1L1 and GFAP in FXS cultures compared to controls further validated this shift in lineage output (**Figure 4D**; **Supplementary Figure 4B**). Together, these findings demonstrate that FXS NPCs follow an abnormal differentiation trajectory, characterized by reduced neuronal commitment and enhanced astrocytic fate acquisition.

### Lowering H3K79me2 levels normalizes proliferation in FXS NPCs

Given the increased proliferation and altered differentiation trajectory observed in FXS NPCs, we next asked whether these phenotypes could be modulated through inhibition of DOT1L, the sole human H3K79 methyltransferase^77^. To this end, CTL and FXS NPCs were treated with two distinct and selective DOT1L inhibitors, EPZ5676 and SGC0946. To confirm the efficacy of DOT1L inhibition, we assessed global H3K79me2 levels after a 72h treatment (**Figure 5A**) and showed a marked reduction of H3K79me2 in both CTL and FXS NPCs following EPZ5676 and SGC0946 treatment compared to vehicle (VHL)-treated conditions (**Figure 5B**). Quantification confirmed an approximately 40% decrease in H3K79me2 levels upon treatment with both inhibitors in CTL and FXS NPCs compared to VHL-treated NPCs (**Figure 5C**; **Supplementary Figure 5**).

**Figure 5.**
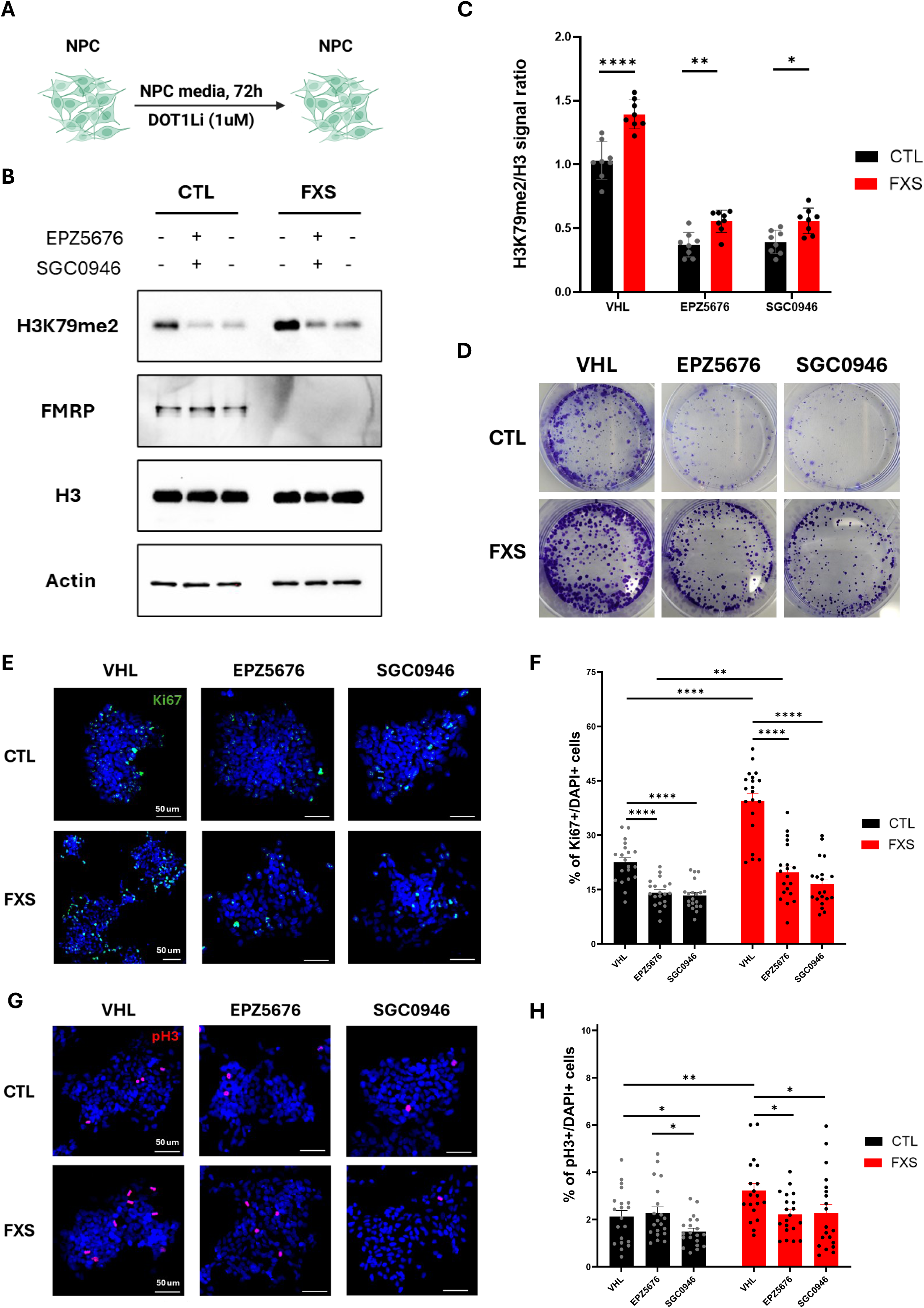
Reducing H3K79me2 levels normalizes proliferation in FXS NPCs. **(A**) NPCs were treated with either 1 μM EPZ5676 or 1 μM SGC0946 for 72h prior to performing experiments. **(B)** Representative western-blot showing reduction of global H3K79me2 levels in NPCs following treatment with DOT1L inhibitors. **(C)** Quantification of H3K79me2 levels obtained by western-blot following treatment with EPZ5676 or SGC0946 (n = 4 cell lines/genotype; 2 replicate per cell line). All the replicates and quantitative analyses can be found in **Supplementary Figure 5. (D)** Representative colony-forming assays of CTL and FXS NPCs following treatment with DOT1L inhibitors. **(E-H)** Representative immunostaining for the proliferation markers Ki67 **(E)** and pH3 **(F)** in CTL and FXS NPCs following treatments with DOT1L inhibitors (top panel), with quantification of Ki67^+^ **(G)** or pH3^+^ **(H)** cells normalized to DAPI^+^ cell counts (n= 4 cell lines/genotype, 5 fields/cell line) (bottom panel). Results are indicated as mean ± SEM. *p< 0.05, ** p<0.01, ***p<0.0001, ****p<0.00001.

Given the on-target activity of both inhibitors, we next investigated whether DOT1L inhibition, leading to reduced H3K79me2 levels, could rescue the proliferation phenotype of FXS NPCs. Colony-forming assays suggest a decrease in the clonogenic potential of FXS NPCs upon treatment with EPZ5676 and SGC0946 (**Figure 5D**; bottom panel). The reduced number and size of colonies formed by FXS cells supported a normalization of their proliferative behavior. Notably, a comparable effect of DOT1L inhibition was observed in CTL NPCs, suggesting that the reduction of H3K79me2 levels is not exclusively associated with correction of the disease phenotype (**Figure 5D**; top panel).

To further validate these observations, we next performed immunofluorescence staining for the proliferation marker Ki67 and showed a significant reduction in the proportion of proliferating cells in FXS NPCs upon treatment with both inhibitors compared to vehicle-treated conditions, as indicated by the percentage of Ki67-positive cells (39 ± 9.5% vs 20 ± 7.8% for EPZ5676 and 17 ± 6.4% for SGC0946, normalized to DAPI+ cells) (**Figures 5E-F**). CTL NPCs population displayed a more modest reduction of proliferating cells upon treatment, indicating that FXS cells are particularly sensitive to DOT1L inhibition. Consistently, immunofluorescence staining for the mitotic marker pH3 revealed a reduced proportion of FXS NPCs in the M phase compared VHL-treated NPCs (**Figures 5G-H**). Together, these results demonstrate that the reduction of H3K79me2 levels by pharmacological inhibition of DOT1L is sufficient to normalize the abnormal proliferative capacity of FXS NPCs.

### DOT1L inhibition restores the differentiation trajectory of FXS NPCs

Considering the impaired neuronal differentiation and altered lineage commitment observed in FXS NPCs, we next examined whether DOT1L inhibition could restore these defects. CTL and FXS NPCs were pre-treated with EPZ5676 and SGC0946 for 72h, followed by differentiation in inhibitor-free conditions (**Supplementary Figure 6A**). Neuronal commitment was then assessed over a 30-day period. Immunofluorescence staining for the neuronal markers MAP2 and TUBB3 at D30 revealed a marked improvement in neuronal differentiation in FXS cultures pretreated with DOT1L inhibitors compared to VHL-treated conditions (**Figure 6A**; right panel). Importantly, DOT1L inhibition also enhanced neuronal lineage marker staining in CTL cultures, indicating that this effect is not restricted to FXS cells (**Figure 6A**; left panel). To further characterize the effect of DOT1L inhibition on differentiation dynamics, we quantified the expression of stage-specific markers over the course of differentiation. Neuronal markers, including NeuN, MAP2, but not GAP43, exhibited enhanced induction in FXS cultures pretreated with DOT1L inhibitors compared to VHL-treated FXS cells, approaching levels observed in VHL-treated CTL NPCs (**Figure 6B**; **Supplementary Figure 6B-D**). In parallel, the downregulation of the NPCs marker PLZF was accelerated upon treatment, suggesting improved exit from the progenitor state and enhanced commitment toward neuronal differentiation (**Figure 6B**; **Supplementary Figure 6E).**

**Figure 6.**
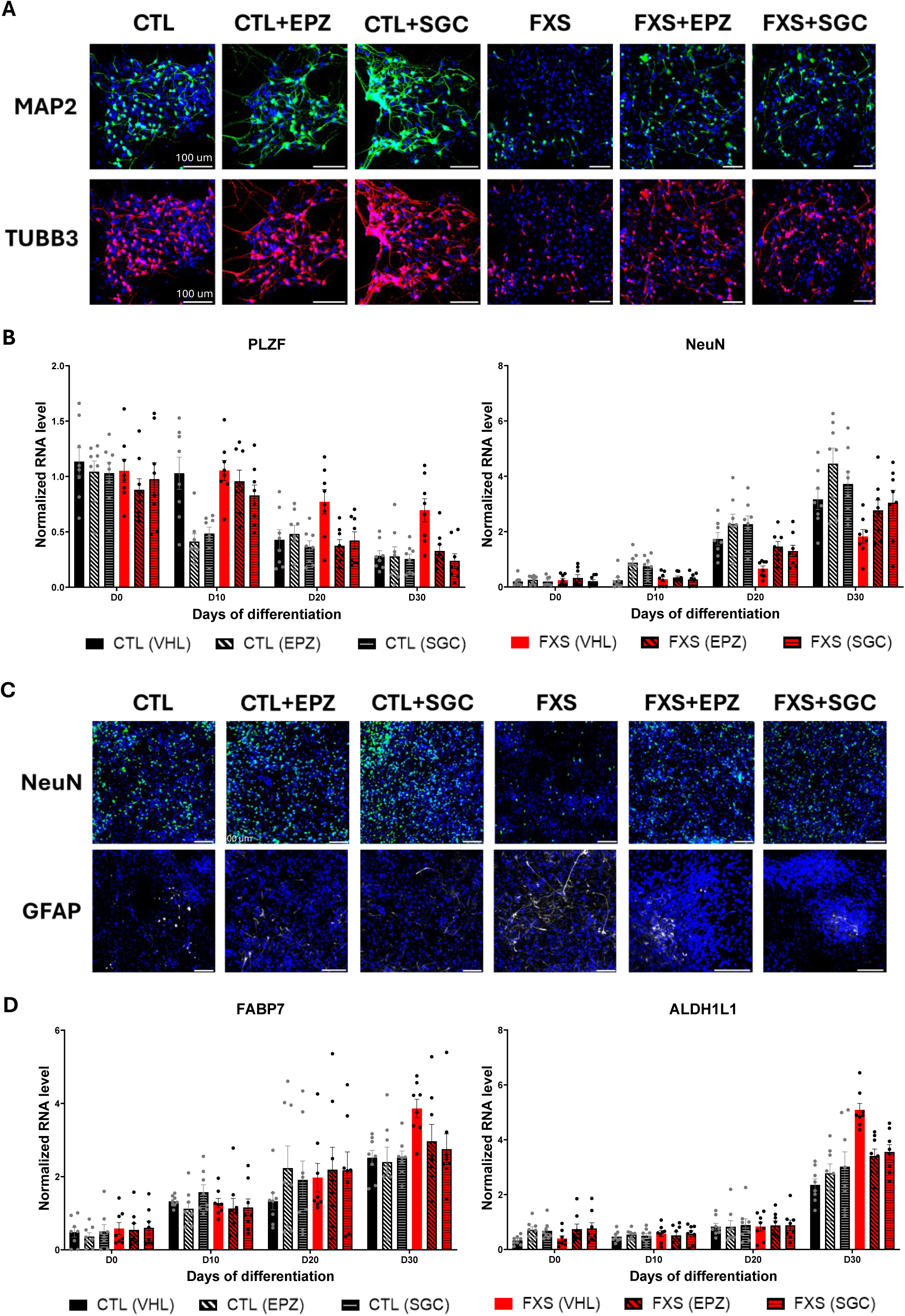
DOT1L inhibition corrects the differentiation trajectory of FXS NPCs. **(A)** Representative immunostaining against the neuronal cytoskeleton markers MAP2 and TUBB3 in CTL and FXS in neural culture at day 30 (D30) following NPC treatment with the DOT1L inhibitors EPZ5676 or SGC0946. **(B)** Quantification of NPC- and neuron-specific markers, respectively PLZF and NeuN, by quantitative PCR during differentiation of CTL and FXS NPCs into neural cultures following NPC with DOT1L inhibitors (n = 4 cell lines/genotype, 2 replicates/cell line). Statistical analyses can be found in **Supplementary Figure 6B-C. (D)** Representative immunostaining against the neuronal marker NeuN and the astrocyte marker GFAP in CTL and FXS in neural culture at day 30 (D30) following NPC treatment with the DOT1L inhibitors EPZ5676 or SGC0946. **(B)** Quantification of astrocyte-specific markers, respectively FABP7 and ALDH1L1, by quantitative PCR during differentiation of CTL and FXS NPCs into neural cultures following NPC with DOT1L inhibitors (n = 4 cell lines/genotype, 2 replicates/cell line). Statistical analyses can be found in **Supplementary Figure 7A-B.**

We next examined whether DOT1L inhibition could correct the lineage bias previously observed in FXS NPCs. Immunofluorescence staining at D30 showed a reduction in the astrocytic marker GFAP in pretreated FXS cultures compared to VHL-conditions, while the expression of the neuronal marker NeuN was concomitantly increased (**Figure 6C**). Consistent with these observations, expression levels of astrocytic markers such as FABP7, ALDH1L1, and GFAP were decreased in treated FXS cultures compared to VHL-treated cells (**Figure 6D**; **Supplementary Figure 7**), further supporting a rebalancing of lineage specification. Together, these results demonstrate that DOT1L inhibition restores the differentiation trajectory of FXS NPCs by promoting neuronal commitment and limiting aberrant astrocytic differentiation, highlighting a key role for DOT1L-dependent epigenetic regulation in cell fate specification.

## Discussion

Our findings revealed that loss of FMRP causes stage-specific dysregulation of global H3K79me2 levels in FXS NPCs, driving widespread transcriptional abnormalities. This altered transcriptomic landscape is defined by reduced expression of neurogenesis-associated genes and elevated expression of programs promoting NPCs proliferation and glial lineage commitment. Functionally, these molecular perturbations manifest as excessive proliferation and a skewed differentiation trajectory, with FXS NPCs displaying diminished neuronal output and a pronounced bias toward astrocytic fate (**Figure 7**).

**Figure 7.**
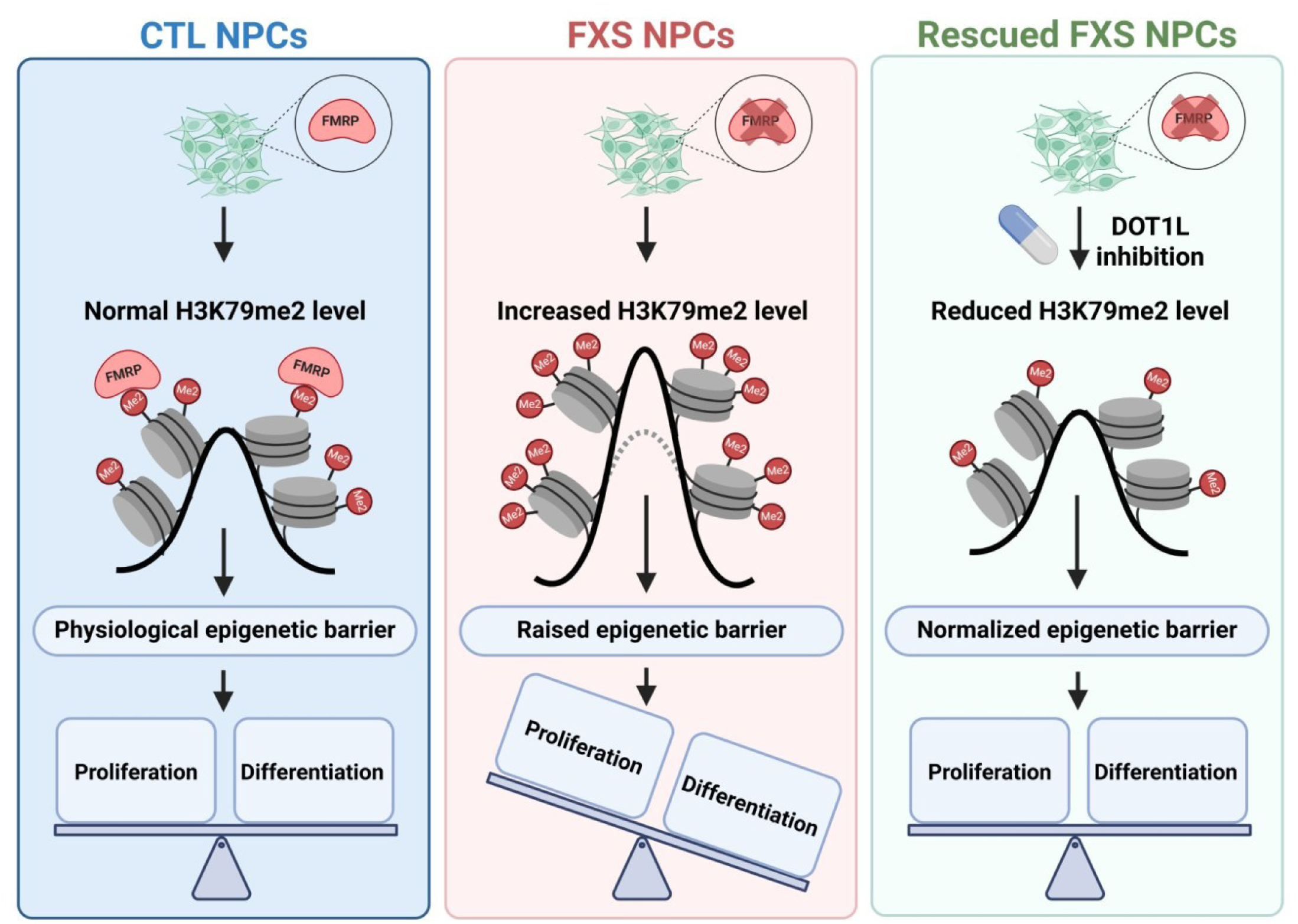
An H3K79me2-dependent epigenetic barrier in neural development. NPCs display physiological levels of H3K79me2, associated with a normal epigenetic barrier and this balanced chromatin state supports proper regulation between proliferation and differentiation (left panel). Loss of FMRP is associated with increased H3K79me2 levels and a strengthened epigenetic barrier. This altered chromatin landscape disrupts normal developmental control, shifting the balance and impairing proper coordination of proliferation and differentiation programs (middle panel). DOT1L inhibition reduces H3K79me2 levels, restoring a more physiological epigenetic state. This normalizes the epigenetic barrier and re-establishes balanced regulation of NPC proliferation and differentiation.

During development, cells transition from pluripotency to lineage-specific states through highly controlled transcriptional programs. A central feature of this process is the establishment of epigenetic barriers. These barriers restrict cellular plasticity by preventing reactivation of pluripotency genes once differentiation has begun, and ensure that committed cells silence alternative developmental trajectories, thereby blocking inappropriate lineage programs^78^. Furthermore, additional studies indicate that these epigenetic barriers also regulate the gradual unfolding of maturation programs during neuronal differentiation^45^. Loss-of-function of specific epigenetic factors in progenitors can lead to premature maturation. In particular, transient inhibition of DOT1L, the only known human H3K79 methyltransferase^77^, at progenitor stage primes cells to acquire mature neuronal properties more rapidly, linking H3K79 methylation to an epigenetic checkpoint controlling neuronal maturation^45^.

Loss of FMRP increases H3K79me2 levels specifically at NPCs stage (**Figure 1B**) and leads to aberrant H3K79me2 occupancy (**Figure 2; Supplementary Figure 2**). One possible mechanism could be that loss of FMRP enhances DOT1L expression. Interestingly, DOT1L mRNA has been identified as an FMRP target in multiple human models, including NPCs^74,79,80^. Given the well-established role of FMRP in regulating the translation of its mRNA targets, it is plausible that the dysregulated H3K79me2 levels observed in FXS NPCs may arise from increased DOT1L expression. Another possible mechanism to explain this dysregulation could stem from the loss of FMRP nuclear functions. Nuclear FMRP could use its RNA-binding capabilities to act as a scaffold, facilitating the recruitment of long non-coding RNAs that could guide epigenetic modifiers, including the H3K79 histone demethylase KDM2B^81^, to specific loci. Loss of FMRP could impair the active demethylation of H3K79me2, contributing to global increase in this epigenetic mark observed in FXS NPCs.

Our findings showed that transient inhibition of DOT1L prior to differentiation is sufficient to markedly improve the proliferation dynamics and the neuronal differentiation capacity of FXS NPCs, with treated cultures reaching levels comparable to those observed in untreated control conditions. This result supports the idea that elevated H3K79me2 levels contribute to the unbalanced proliferation and differentiation trajectory of FXS progenitors (**Figures 3 and 4**) and that their reduction can restore these phenotypes (**Figures 5 and 6)**. The ability to normalize neuronal output following a relatively short pre-treatment further suggests that epigenetic dysregulation in FXS NPCs remains reversible and that early modulation of chromatin state can have lasting effects on cell fate decisions. Notably, DOT1L inhibition also reduced proliferation and promoted neuronal marker expression in control NPCs, indicating that its effects are not limited to the disease context. Rather than functioning exclusively as a corrective mechanism, reduced H3K79me2 levels may more broadly enhance the competence of NPCs to undergo neuronal differentiation. These findings suggest that DOT1L activity participates in the regulation of a general epigenetic barrier that influences the timing or efficiency of neuronal commitment. Within this framework, FXS NPCs may display an exacerbated form of this barrier, making them particularly responsive to modulation by DOT1L inhibitors.

Our results also indicate that FXS NPCs display a biased differentiation trajectory, preferentially committing to astrocytic lineage at the expense of neuronal output. Consistent with this, a recent study reported increased astrocytes density in the cortex of FXS patients^82^. Astrocytes play critical roles in supporting neuronal functions, including synapse formation and maturation, as well as the regulation of synaptic transmission^83,84^. Therefore, an altered neuron-to-astrocyte ratio in FXS could underlie key neurophysiological abnormalities observed in patients, such as impaired dendritic spine morphogenesis, disrupted excitatory–inhibitory balance, and weakened neuronal connectivity^85^. These insights highlight the potential of therapeutic strategies that target astrocytic function or correct NPC lineage bias as promising avenues for the treatment of FXS.

Taken together, our findings support the existence of an H3K79me2-dependent epigenetic barrier that plays a central role in regulating NPCs fate decisions. Disruption of this barrier in FXS leads to altered proliferation dynamics and impaired differentiation, highlighting its importance in maintaining proper developmental trajectories. Future studies should aim to further characterize the molecular mechanisms governing this epigenetic barrier and how its modulation influences neural lineage commitment. Importantly, FMRP has also been shown to associate with other histone modifications, including H3K27me1 and H3K36me3^22^, both of which are enriched at actively transcribed genes^86,87^. Investigating how these marks contribute to the establishment or regulation of this epigenetic barrier, and whether their dysregulation participates in FXS pathophysiology, will be essential to refine our understanding of the epigenetic control of neural development.

## Acknowledgments

The authors thank Edouard W. Khandjian for generously providing the anti-FMRP IgY antibody and for his valuable insights on this manuscript. The authors thank Dr. Silvia Di Angelantonio and Dr. Alessandro Rosa from Sapienza University for providing the IPS_WT and IPS_KO cell lines. The authors thank Dr Kagistia Hana Utami for kindly providing help with the differentiation protocols. This work was supported by a grant from the Canadian Institutes of Health Research (CIHR) to B.L. [PJT-166109]. This work was also supported by grants from the CHUS Research Center (Sherbrooke, Canada) and the FRAXA Foundation to F.C., B.L and O.D.

O.D. holds doctoral scholarships from CIHR, Fonds de Recherche du Québec - Santé (FRQS), Brain Canada Foundation and the Corporation for Research and Action on Hereditary Diseases. G-L.G. was supported by an Undergraduate Research Award (URA) from the Natural Sciences and Engineering Research Council of Canada (NSERC). C.B-S. was supported by a post-doc fellowship from the Research Center on Aging (CdRV, Sherbrooke, Canada).

## Material and Methods

### iPSC culture and differentiation

Four iPSC lines derived from healthy individuals (IPS_C1, IPS_C2, IPS_C3 and IPS_WT)^49,51^, three iPSC lines derived from FXS individuals (IPS_X1, IPS_X2, IPS_X3)^49^ and one FMR1 Knock-out (isogenic to IPS_WT) iPSC line (IPS_KO)^50^ were used in this study. Please refer to **Supplementary table 1** for a more extensive description of these cell lines.

All iPSC lines were cultured on Matrigel-coated plates in mTeSR1 medium with daily medium change. Single-cell passaging was performed when cells reached a confluency of 80%. Briefly, cells were washed once with dPBS and treated with Accutase for 5 min at 37°C. Cells were then harvested with dPBS, centrifuged at 400g for 5 min, and seeded on Matrigel-coated plates in mTeSR1 medium supplemented with 10µM Y-27632 using a 1:10 splitting ratio. For cryopreservation, single-cell suspension of iPSC were resuspended in mFreSR and frozen using a standard slow-cooling procedure^88^.

iPSCs were differentiated into neural progenitor cells (NPCs) using an adaption of previously published protocols^47,89^. iPSCs were dissociated into a single-cell suspension as described above and seeded on Matrigel-coated plates in NPC medium (NPCM) supplemented with 0.1µM compound E and 10µM Y-27632 at a density of 30,000 cells/cm^2^. Culture medium was changed everyday with NPCM + 0.1µM compound E for 6 days. On day 7 (D7), NPCs were harvested using Accutase as described above and seeded on Matrigel-coated plates in NPCM + 10µM Y-27632 using 1:4 splitting ratio. Culture medium was changed daily with NPCM. Cells were passaged upon reaching a confluency of 85% with Accutase using a 1:4 splitting ratio. For cryopreservation, single-cell suspensions of NPCs were resuspended in NPCM + 10% DMSO and frozen using a standard slow-cooling procedure. NPCM was composed of: DMEM/F12: Neurobasal (1 :1), 1% N2-A, 2% B-27 (without retinoic acid), 1% PenStrep, 1% GlutaMAX, 10ng/ml hLIF, 5ug/mL BSA, 4uM CHIR99021, 3 mM SB431542.

NPCs were differentiated into mature neural culture using an adaptation of previously published protocols^48,89^. Briefly, NPC were harvested using Accutase as described above and seeded on 40µg/ml poly-L-ornithine + 20µg/ml laminin-coated plates in neural differentiation medium (NeuroDiff) supplemented with 10µM Y-27632 at a density of 20 000 cells/cm^2^. The following day, the culture medium was replaced with NeuroDiff and subsequently refreshed every 2–3 days. NeuroDiff was composed of: DMEM/F12: Neurobasal (1 :1), 1% N2-A, 2% B-27, 1%, 20 ng/ml BDNF, 20ng/ml GDNF, 200 nM L-ascorbic acid, 300 uM dibutyryl-cyclic AMP.

To modulate the H3K79me2-dependent epigenetic barrier, NPCs were treated with 1 μM of EPZ5676 or SGC0946 DOT1L inhibitors for 72 hours, with daily medium replacement. After treatment, cells were either collected for downstream analyses or subjected to differentiation without DOT1L inhibitors, as described above.

### Whole cell protein extraction and quantification

Cells were washed once with ice-cold dPBS before being lysed by applying ice-cold RIPA buffer directly into the culture dish. RIPA buffer was composed of 50mM Tris-HCl pH 8.0, 150mM NaCl, 1%NP-40, 0.5% sodium deoxycholate,0.1% SDS and was supplemented with 1% P8340 protease inhibitor cocktail and 1% halt phosphatase inhibitor cocktail immediately before use. Cell extracts were harvested using a cell scrapper, treated with 500 units of Benzonase Nuclease (10 min, 4°C) and spun at 20,000g (4°C) for 10 min. The resulting supernatant were transferred into a clean microcentrifuge tube and stored at -80°C. Protein quantification was performed using bicinchoninic acid assay (BCA).

### Whole cell RNA extraction and quantification

Cells were washed once with ice-cold dPBS before being lysed by applying TRIzol reagent directly into the culture dish. Cell extracts were harvested using a cell scrapper and spun at 20,000g (RT) for 10 min. The resulting supernatant was transferred to a clean microcentrifuge tube and total RNAs were extracted using the Direct-zol RNA miniprep kit following the manufacturer’s instructions. RNA quantification was performed using a NanoDrop spectrophotometer (ThermoFisher Scientific).

### Western Blot

Protein samples were boiled in 1X Lammeli buffer (60mM Tris-HCl pH 6.8, 2% SDS, 10% glycerol, 5% 2-mercaptoethanol and 0.5% bromophenol blue) for 5 min before being resolved on SDS polyacrylamide gels. Proteins were transferred on nitrocellulose membrane (0.45µM, Biorad) and block using 5% non-fat dry milk in tris buffer saline (TBS) with 0.1% Tween-20 (TBS-T) for 60 min at room temperature. Primary antibodies were diluted in 5% non-fat dry milk in TBS-T supplemented with 0.02% sodium azide and hybridized by overnight incubation at 4°C. The following primary antibodies were used: FMRP (1:1000, Millipore Sgima #MAB2160), beta-Actin (1:5000, Cell Signaling Technology #4970), phospho-AKT (1:2000, Cell Signaling Technology #4060), AKT (1:2000, Cell Signaling Technology #2920), phospho-ERK (1:2000, Cell Signaling Technology #4370), ERK (1:2000, Cell Signaling Technology #4696), γH2AX (1:2000, Cell Signaling Technology #9718), H3 (1:5000, Cell Signaling Technology #9715), H3K79me2 (1:2000, Active Motif #39143), alpha-Tubulin (1:2500, Abclonal #A6830), Puromycin (1:500 DSHB #PMY-2A4-s). Membranes were then washed three times with TBS-T and incubated with secondary antibodies (diluted in 5% non-fat dry milk in TBS-T) for 60 min at room temperature. The following secondary antibodies were used: anti-mouse HRP (1:10 000, Jackson ImmunoResearch #715-035-150), anti-rabbit HRP (1:10 000, Jackson ImmunoResearch #711-035-152). Immunoblots were revealed using Clarity Western ECL substrate (Biorad #1705061) or Clarity Max Western ECL substrate (Biorad #1705062) and imaged using an iBright FL1500 (ThermoFisher Scientific). Quantification was performed using the Fiji software (version 2.16.0)^90^.

### Immunofluorescence

Cells were grown on Matrigel or PLO/laminin-coated glass coverslips and fixed with 4% paraformaldehyde (diluted in the culture media) for 20 min at 37°C. Cells were then washed three times with dBPS and incubated with blocking solution (10% normal goat serum (NGS), 0.1% Triton X-100 in dPBS) for 60 min at room temperature. Primary antibodies were diluted in 10% NGS (diluted in dPBS) and incubated with the coverslips overnight at 4°C. The following primary antibodies were used: SOX2 (1:100, SantaCruz Biotechnology #sc-365823), OCT4 (1:250, Cell Signaling Technology #2840), Nanog (1:500, Abclonal #A3232), FMRP (1:200, Dury et al., 2013^91^), PAX6 (1:500, Biolegend #901301), Nestin (1:100, Santa Cruz Biotechnology #sc23927), H3K79me2 (1:250, Active Motif #39143), MAP2 (1:250, Cell Signaling Technology #4542), TUBB3 (1:250, R&D systems #MAB1196), GFAP (1:250, Cell Signaling Technology #3670), NEUN (1:250, Cell Signaling Technology #24307), phospho-H3 (1:250, Cell Signaling Technology #9706) and Ki67 (1:250, Cell Signaling Technology #9129). Coverslips were washed three times with dPBS and incubated with secondary antibodies (diluted in 10% NGS) for 60 min at room temperature protected form light. The following secondary antibodies were used: Anti-Rabbit IgG 488nm (1:1000, Cell Signaling Technology #4412), Anti-mouse IgG 647nm (1:1000, Cell Signaling Technology #4410), Anti-chicken IgY 647nm (1:1000, Jackson ImmunoResearch #103-605-155). Coverslips were washes three times and counterstain with DAPI (1µM in dPBS, 10 min at room temperature). Coverslips were then mounted on microscopic slides using SlowFade Gold Antifade Mountant (ThermoFisher) and sealed with nail polish. Slides were observed using either an epifluorescence microscope (Zeis AxioSkop2, Carl Zeiss, Inc., Thornwood, NY) with a 10X or 40X objectives or with a confocal microscope (Olympus FV100, Olympus, Tokyo, Japan) using an 60X oil immersion objective. Images were analyzed using the Fiji software (version 2.16.0)^90^.

### Reverse transcription and quantitative PCR

Up to 1000 ng of total RNA were used to generate cDNA using iScript cDNA synthesis kit (Biorad #1708891) following instructions from the manufacturer. Quantitative PCR (qPCR) reactions were performed with the PerfeCta SYBR Green SuperMIx (QuantaBio #95054) using 5 ng of cDNA on an Azur Cielo 3 real-time PCR system (Azure Biosystems, Dublin, CA, USA). Real-time PCR reactions were performed with a hot start step at 95°C for 2 min, followed by 40 cycles of 95⁰C for 10 seconds, 60⁰ for 10 seconds and 72°C for 20 seconds. Analysis was performed using the Azur Cielo Manager software (Azure Biosystems) and gene expression normalized to ACTB levels. Primers used in this study are listed in **Supplementary Table 2**.

### Proliferation curves

Single-cell suspensions of NPCs were seeded on Matrigel-coated 12-well plates (10,000 cells/well) in NPCM + 10µM Y-27632. Culture medium was refreshed daily with NPCM. Starting 2 days following plating (D3), cells from three wells were harvested everyday until day 7 using Accutase and viable cells were counted using Trypan Blue exclusion on an hemacytometer.

### Colony forming assay

Single-cell suspensions of NPCs were seeded on Matrigel-coated 6-well plates (5,000 cells/well) in NPCM +10 µM Y-27632. Culture media was refresh daily with NPCM for 9 days. On D9, cells were washed once with dPBS and fixed by incubating with 4% paraformaldehyde (diluted in NCPM) 20 min at 37°C. Cells were washed again three times with dPBS and stain with 0.5% Crystal violet for 60 min at room temperature. Cells were washed multiple times with ddH_2_O to remove excess stain and air dry for 2 hours and photography of the well were taken using a digital camera.

### Cell cycle profiling

Cell cycle profiling was performed through quantification of DNA content by propidium iodide (PI) staining. In summary, NPC were harvested with Accutase and pelleted by centrifugation (400g, 5 min). The pellet was resuspended in 0.5 ml dPBS, and cells were fixed with 3 ml cold (-20°C) 70% ethanol added dropwise. NPCs were then centrifuged at 400g for 10 min, washed two time with ice-cold dPBS and incubated with the staining solution (dPBS, 0.1% Triron-X100, 5µg/ml PI and 50µg/ml RNAse A) 30 min at 4°C protected from light. Cells were analysed on a CytoFLEX 20 flow cytometer (Beckman Coulter Life Sciences) equipped with an automatic 96 wells plate loader and a 488nm laser. Data acquisition was set to collect a minimum of 100,000 single-cells per sample and flow rate was set to 15µL/min. Forward scatter area (FSC-A); side scattered area (SSC-A) and SSC-width signals were used to establish the live gates to exclude debris, and cell clumps. Data were acquired using the CytExpert software (v2.5, Beckman Coulter Life Sciences) and cell cycle profiling analysis performed with ModFit LT (v6.0, Verity Software House).

### Cell synchronization

NPC were synchronized at different phases of the cell cycle trough 16 hours treatment with small molecules. Synchronization at the G1 phase was achieved with 1µM PD-0332991^76^, G2/M border with 10µM Ro-3306^75^ or with 100ng/ml nocodazole^71^. Synchronization efficacy was validated by cell cycle profiling as described above (**Supplementary Figure 3**).

### Cell fractionation

The cell fractionation procedure used in this study was adapted from a previously published protocol^92^. In summary, cells were harvested with Accutase and pelleted by centrifugation (400g, 5 min) and washed once with ice-cold dPBS. The cell pellet was then resuspended in cold cell lysis buffer (5mM PIPES pH8.0, 80mM KCl, 0.5% NP-40 and 1% P8340 supplemented immediately before use), incubated on ice for 10 min and centrifuged at 800 for 5 min (4°C). The resulting supernatant (containing the cytoplasmic, membrane and mitochondrial fractions) was transferred into a new tube and the pellet (containing the nuclei) was washed once with cold cell lysis buffer. The nuclei pellet was then resuspended in cold nuclear lysis buffer (20 mM Tris-HCl pH 7.4, 400 mM NaCl, 0.5% NaDOC, 0.1%SDS and 1% P8340 supplemented immediately before use) and stored at -80°C.

### FMRP immunoprecipitation (IP)

A previously described protocol was adapted to immunoprecipitate FMRP from nuclear fractions^93^. Lysate containing the nuclear fractions were treated with Benzonase nuclease (750 units) for 15 min on ice and then centrifuged at 20,000g for 10 min (4°C). The collected supernatant was raised to 10 mM EDTA. For each IP reaction, 100ug of protein from nuclear fraction was incubated with 50µg of anti-FMRP IgY antibody^91^ or 50µg of non-immunized IgY immobilized on 50µl of anti-chicken IgY agarose beads. Samples were incubated overnight at 4°C under rotation and washed four times with IP wash buffer (20 mM Tris-HCl pH 7.4, 400 mM NaCl, 0.5% NP-40 and 10 mM EDTA). Immunoprecipitated proteins were eluted by the addition of 2X Lammeli buffer (120mM Tris-HCl pH 6.8, 4% SDS, 20% glycerol, 10% 2-mercaptoethanol and 1% bromophenol blue), boiled (95°C) for 5 min and analyzed by Western Blot.

### Chromatin immunoprecipitation (ChIP)

Cells (5x10^6^) were crosslinked with 1% formaldehyde for 10 min at room temperature. The reaction was terminated by the addition of 125 mM glycine (5 min at room temperature). Cells were then wash with dPBS, harvested with a cell scrapper and centrifuged at 400g for 5 min. Cells were resuspended in lysis buffer (50 mM Tris pH8.0,10 mM EDTA pH8.0, 1% SDS, 1% P8340) and DNA was sheared into 300-600 base-pair fragments trough sonication using a Sonic Dismembrator Model 500 (ThermoFisher Scientific) at 15% amplitude, applying a cycle of 10 seconds on and 40 seconds off, for a total sonication duration of 4 min. SDS concentration in sheared chromatin samples was reduced to 0.1% with the addition of 9 volumes of dilution buffer (16.7 mM Tris-HCl pH 8, 0.01% SDS, 1.1% Triton X-100, 1.2 mM EDTA pH 8.0, 167 mM NaCl, 1% P8340). Chromatin fragments (30% of the whole sample) were incubated overnight at 4°C with 9µg H3K79me2 antibody (Active Motif #39143) or with 9µg normal rabbit IgG (Cell Signaling Technology #2729) and 40µl of pre-washed Dynabeads Protein A (ThermoFisher Scientific, #10002D). Beads were sequentially washed with the following buffers: Once (4°C) with low salt wash buffer (20 mM Tris-HCl pH 8, 0.1% SDS, 1% Triton X-100, 2 mM EDTA,150 mM NaCl), once (4°C) with high salt wash buffer (20 mM Tris-HCl pH 8, 0.1% SDS, 1% Triton X-100, 2 mM EDTA, 500 mM NaCl), once (4°C) with LiCl wash buffer (10 mM Tris pH 8,0.25 M LiCl,1% NP-40,1% NaDoc,1 mM EDTA) and twice (room temperature) with TE buffer (10 mM Tris-HCl,1 mM EDTA). DNA was eluted twice with 200µl Elution buffer (100mM NaHCO3, 1% SDS) incubated at 65°C for 15 min. The combined eluates reversed cross-link with 200 mM NaCl (overnight at 65°C) and sequentially treated with 10ug RNAse A (37°C, 30 min) and 40µg proteinase K (50°C, 3 hours). DNA was then recovered by phenol/chloroform extraction and precipitated overnight at -20°C in the presence of 70% ethanol, 100 mM sodium acetate and 20µg/ml glycogen. Samples were resuspended in 10µl of water and quantified using the Qubit 1X dsDNA HS Assay kit (ThermoFisher) on a Qubit spectrofluorometer (ThermoFisher).

### DNA sequencing following chromatin immunoprecipitation

ChIP-seq libraries were prepared using the NEBNext Ultra II DNA Library Prep Kit (New England Biolabs). Libraries were prepared using 20ng of immunoprecipitated DNA following the manufacturer’s instructions. Each library was purified and size-selected (200-700 bp) using SPRIselect beads (Beckman Coulter) and quantified using the Qubit 1X dsDNA HS Assay kit (ThermoFisher) on a Qubit spectrofluorometer (ThermoFisher). A multiplex library was prepared by pooling an equimolar volume of each individual library and concentrated using SPRIselect beads (Beckman Coulter). The resulting library was then sequenced at RNomics platform of the Université de Sherbrooke (Sherbrooke, Canada). Single-end sequencing was performed on all samples with a 75 bp sequencing depth on a Aviti sequencer (Element Biosciences).

Quality control, quality and adapter trimming of sequencing libraries were performed with Trim Galore. Reads were aligned with bwa mem against human genome version GRCh38 in single-end mode. Resulting SAM files to BAM file conversion and mapping results were performed with samtools. The following steps were performed in Galaxy environment for computational memory and storage purposes: sorting BAM files (with samtools), removing duplicate molecules (with picard) and BAM filtering on MAPQ quality score >= 20 (with ngsutils). Broad peaks were called with epic2. Resulting BED files were annotated, compared and visualized with R package ChIPseeker. Differential peaks abundance analysis was performed in the Perseus software using a two-tailed student T-test. Permutation-based adjusted p-value method (significance threshold of < 0.05) was conducted to account for multiple testing problem. Peak profile heatmaps were generated in Galaxy with deeptools. Bigwig files for visualization and storage purposes were generated with bedGraphToBigWig. The raw data have been deposited at the Gene Expression Omnibus (GEO) under the subseries entry GSE309276.

### RNA-seq preparation and analysis

RNA-seq libraries were prepared using NEBNext Single Cell/low Input RNA library preparation kit (New England Biolabs). Libraries were prepared using 10ng of total RNA following the manufacturer instructions until the final library purification. Each final library was purified with the MinElute PCR purification kit (Qiagen) and then quantified using the Qubit 1X dsDNA HS Assay kit (ThermoFisher) on a Qubit spectrofluorometer (ThermoFisher). A multiplex library was then prepared by pooling equimolar volumes of individual libraries and size-selected using SparQ PureMag Beads (Quantabio). The resulting library (300-700 base pairs) was then sequenced at RNomics platform of the Université de Sherbrooke (Sherbrooke, Canada). Single-end sequencing was performed on all samples with a 75 bp sequencing depth on a Aviti sequencer (Element Biosciences).

Paired-end fastq quality control, trimming and adpter-removal were performed with Trim Galore. Reads were aligned against human genome version GRCh38 and annotated with gtf version 108 with R package Rsubread. Resulting ensembl gene annotation was matched with gene symbol, description and entrezid by using R package org.Hs.eg.db. Unsupervised MDS and heatmap plots were generated to assess the quality of sample libraries. Library size normalization (through TMM method) and lowly expressed gene filtering were performed using R package edgeR. Differential expression analysis was performed using the Perseus software using a two-tailed student T-test. Permutation-based adjusted p-value method (significance threshold of < 0.01) was conducted to account for multiple testing problem. Gene ontology enrichment analysis were performed using ShinyGO version 0.82^94^. The raw data have been deposited at the Gene Expression Omnibus (GEO) under the subseries entry GSE309275.

### Statistical analysis

Unless specified otherwise, data is represented by mean ± SEM and statistical significance for the difference between control (CTL) and FXS groups was determined using two-tailed student-test conducted in Prism (significance threshold < 0.05). One-way ANOVAs conducted in Prism were performed to determine statistical significance between groups for RT-qPCR data obtained following NPC differentiation after treatments with DOT1L inhibitors. Pearson correlation analyses and Fisher’s exact test were also performed with Prism.

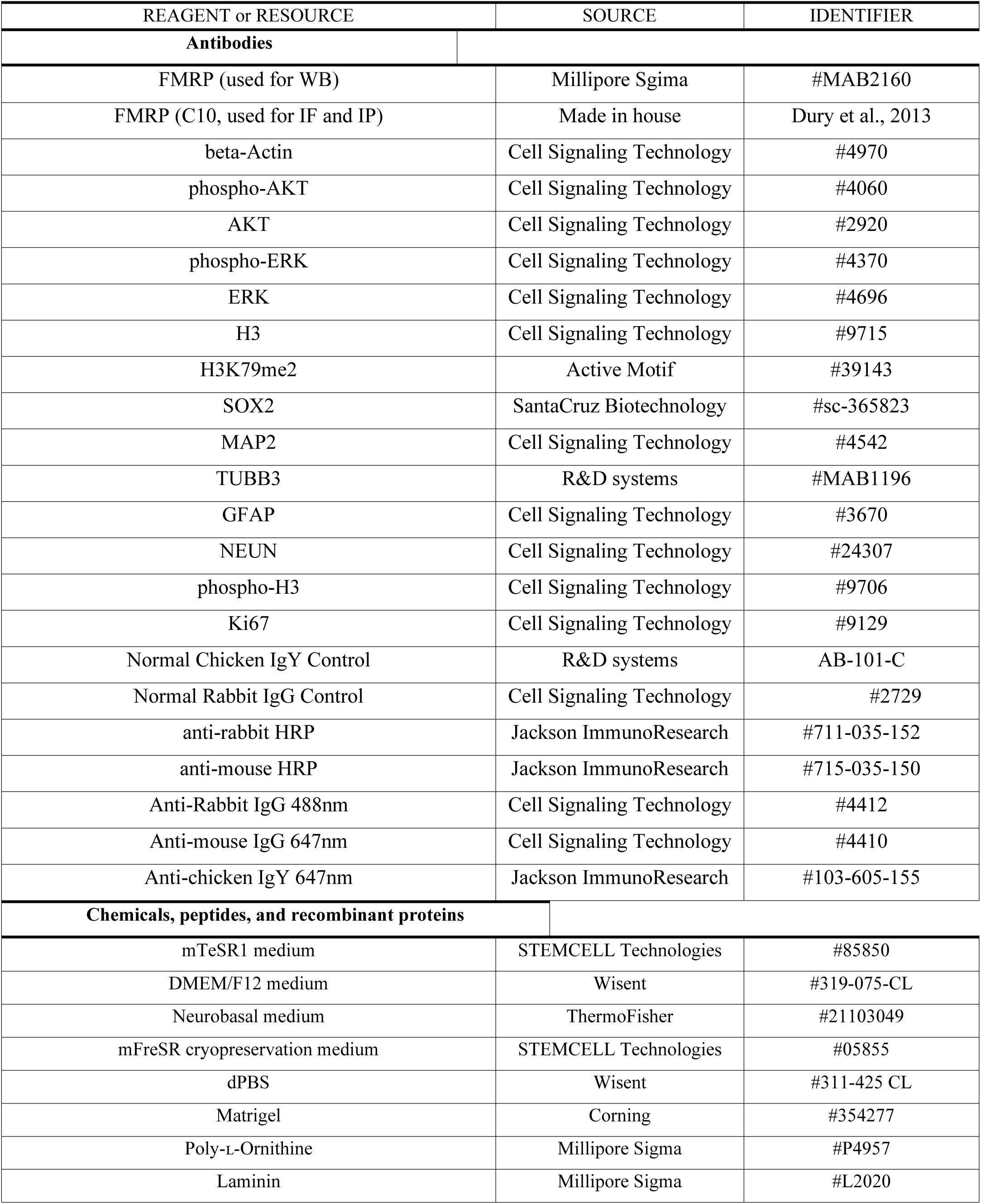
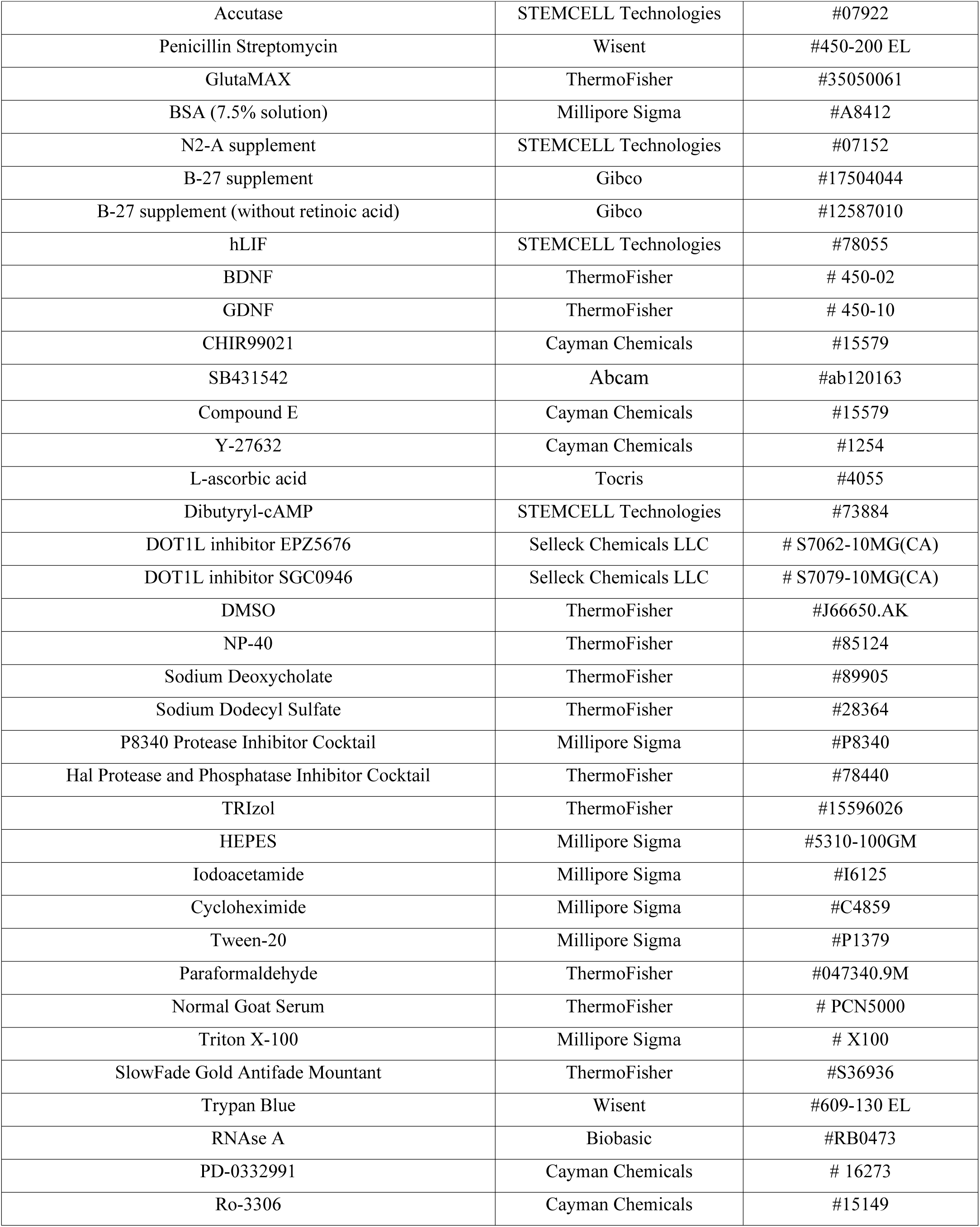
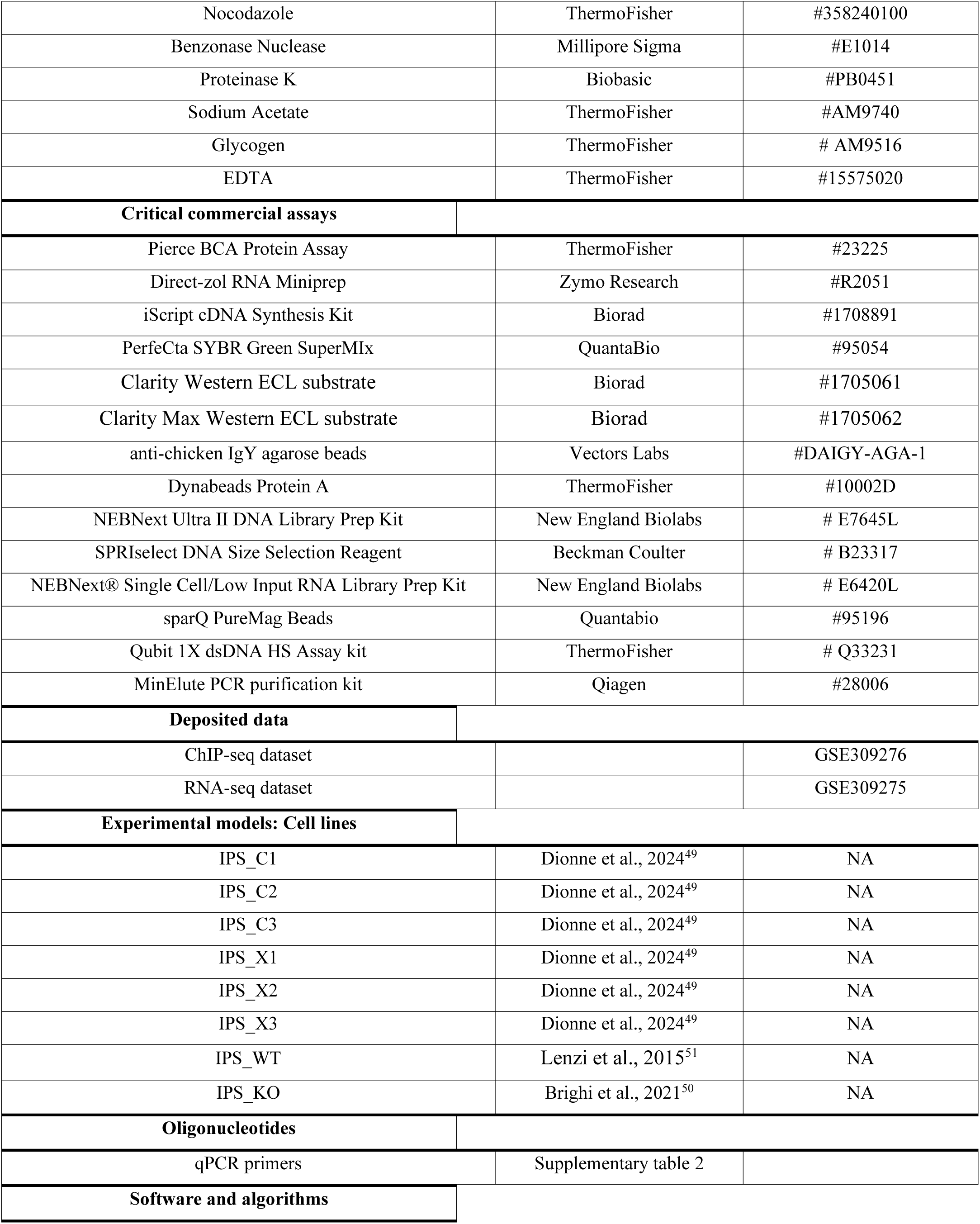
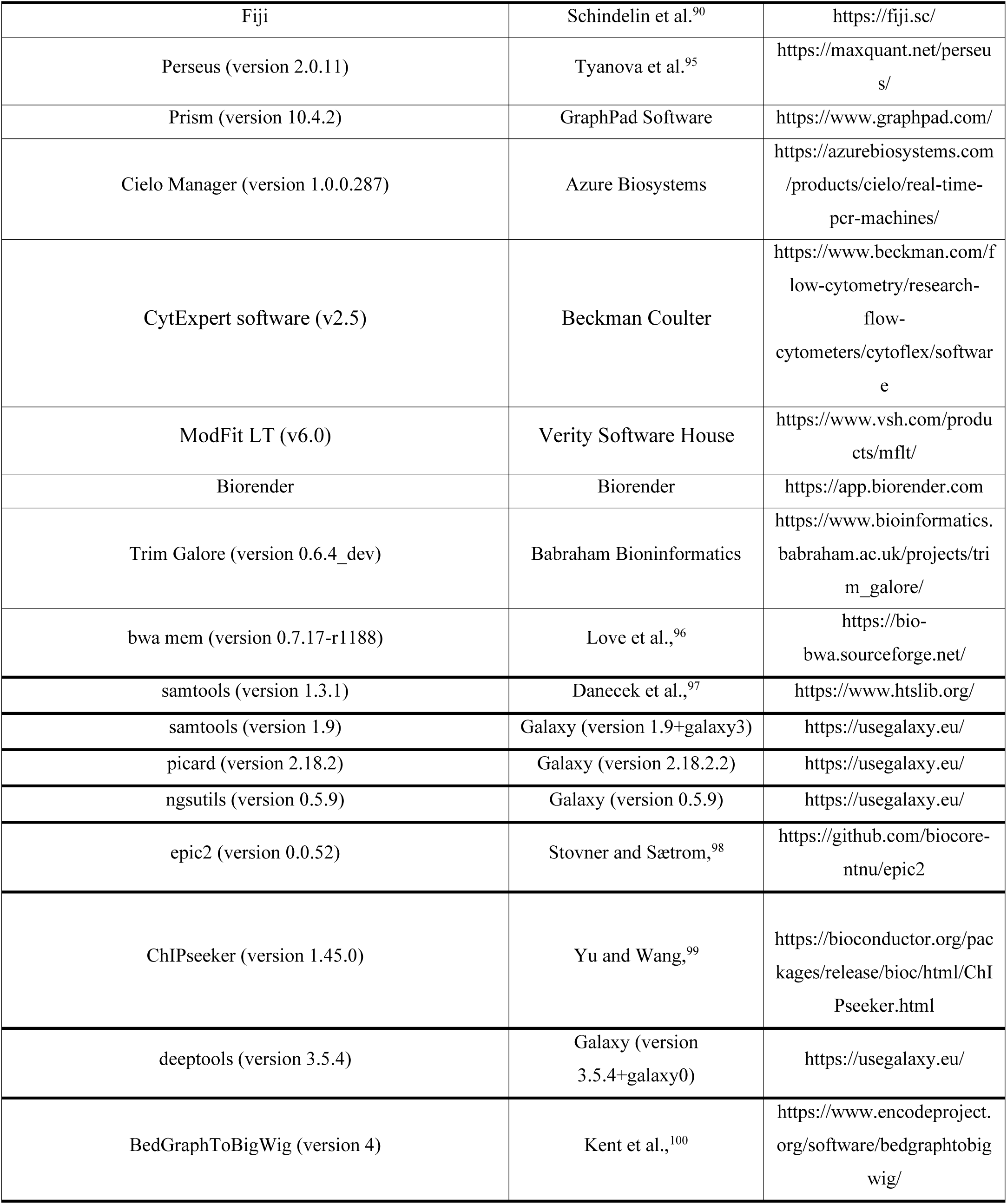
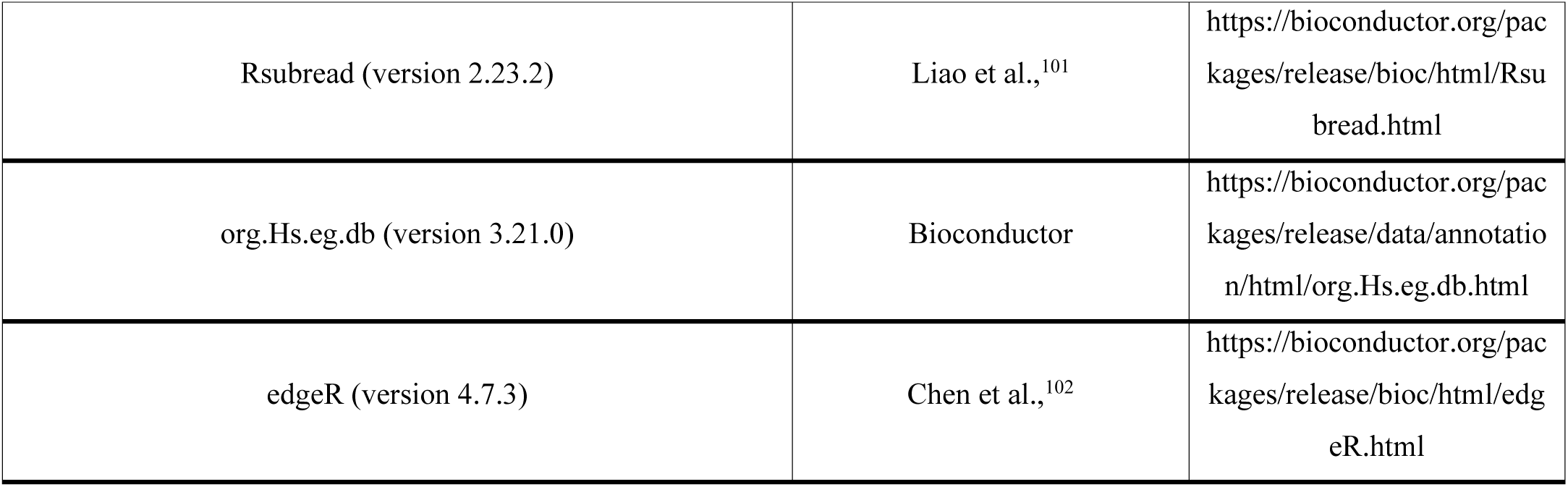

