## Supplemental material for "Dysregulation of an H3K79me2-dependent epigenetic barrier impairs neural progenitor cell proliferation and differentiation in Fragile X syndrome"

### Supplemental tables

### Supplemental table 1. Induced pluripotent stem cell lines used in this study

| Cell line | Sex | Age | Type | Modifications | Primary cells | Reprograming method | Reference |
|---|---|---|---|---|---|---|---|
| IPS_C1 | M | 27 | CTL | NA | Urine-derived cells | Sendai (Oct4, KLF4, Sox2, Myc) | Dionne et al., Front. Cell Dev. Biol (2024) |
| IPS_C2 | F | 22 | CTL | NA | Urine-derived cells | Sendai (Oct4, KLF4, Sox2, Myc) | Dionne et al., Front. Cell Dev. Biol (2024) |
| IPS_C3 | M | 14 | CTL | NA | Urine-derived cells | Sendai (Oct4, KLF4, Sox2, Myc) | Dionne et al., Front. Cell Dev. Biol (2024) |
| IPS_X1 | M | 12 | FXS | NA | Urine-derived cells | Sendai (Oct4, KLF4, Sox2, Myc) | Dionne et al., Front. Cell Dev. Biol (2024) |
| IPS_X2 | F | 29 | FXS | NA | Urine-derived cells | Sendai (Oct4, KLF4, Sox2, Myc) | Dionne et al., Front. Cell Dev. Biol (2024) |
| IPS_X3 | F | 23 | FXS | NA | Urine-derived cells | Sendai (Oct4, KLF4, Sox2, Myc) | Dionne et al., Front. Cell Dev. Biol (2024) |
| IPS_WT | M | 8 | CTL | NA | Fibroblast | Lentivirus (Oct4, KLF4, Sox2, Myc) | Lenzi et al., Dis. Model & Mech., (2015) |
| IPS_KO | M | 8 | FXS | FMR1-KO | Fibroblast | Lentivirus (Oct4, KLF4, Sox2, Myc) | Brighi et al., Cell Death and Disease (2021) |

F = female; M = male; CTL = control; FXS = fragile X syndrome; NA =non-applicable

**Supplemental table 2. List of primers used quantitative PCR**

| Primer | Sequence |
|---|---|
| GAP43-f | GAAGAAGGCAAGGGACGAGACAACC |
| GAP43-r | CTCAGCAGCTTGGACATCATCCTTC |
| NeuN-f | GTCGTGTATCAGGATGGATTTTATGGTG |
| NeuN-r | ATGGTTCCAATGCTGTAGGTCGCC |
| MAP2-f | GGCGGACGTGTGAAAATTGAGAGTGTAA |
| MAP2-r | ACGCTGGATCTGCCTGGGGAC |
| PLZF-f | TGTGGGGTCGAGCTTCCTGATAACG |
| PLZF-r | GCATCCTCCTTCGAAAACTGTGCACC |
| ALDH1L1-f | CAGACAAGGATGGAAAGGCCGACC |
| ALDH1L1-r | CCATGCCGGGGGGCACTGATTA |
| GFAP-f | CGCCAGGATCCCCGCTTTGAAATCTG |
| GFAP-r | TCGGCCGCCAGCTCTTTGATGTGTTC |
| FABP7-f | AGTCAGAACTTTGATGAGTACATGAAGGCTCT |
| FAB7-r | ATCTGCAGTGGTTTCATCAAACTCTTCTCC |
| ACTB-f | GCGGGAAATCGTGCGTGACATT |
| ACTB-r | CTAGAAGCATTTGCGGTGGA |

**Supplementary figure legends**

**Supplementary Figure 1**

**(A-B)** Control **(A)** and FXS **(B)** induced pluripotent stem cells (iPSCs) were differentiated into neural progenitor cells (NPCs) and mature neural culture (neuro). Protein levels of H3K79me2 across the different stages of the differentiation were analyzed by western-blot (left panel). On the right panel, quantification of H3K79me2 global levels (n =4/genotype). Results are indicated as mean ± SEM. **(C)** Immunoblots of FMRP co-immunoprecipitation with H3K79me2 within Benzonase nuclease-treated nuclear fractions of NPCs derived from in the IPS_C1 (NPC_C1), IPS_C1 (NPC_C2) and IPS_WT (NPC_WT) cell lines.

**Supplementary Figure 2**

**(A)** High-resolution profiles of H3K79me2 distribution across PAX6, DLL1, WNT7A and FGFR2 genes, genes involved in the regulation of NPC proliferation and differentiation. **(B)** RNA expression levels of PAX6, DLL1, WNT7A and FGFR2 genes quantified by RNA-seq. **(C-D)** High-resolution profiles of H3K79me2 distribution across genes involved in **(C)** neurogenesis (SLIT1, PIEZO1) and **(D)** astrogenesis (SOX8, SOX9). **(E-F)** RNA expression levels of SLIT1, PIEZO1, SOX8, and SOX9 genes quantified by RNA-seq (n= 4 cell lines/genotype). Results are indicated as mean ± SEM. $*p < 0.05$, $** p<0.01$, $***p<0.0001$, $****p<0.00001$.

**Supplementary Figure 3**

**(A)** Representative immunostaining of phosphorylated histone H3 at serine 10 (pH3) with quantification of $pH3^+$ cells normalized to $DAPI^+$ cell counts (n = 4 cell lines/genotype, 5 fields/cell line). **(B)** Validation of synchronization efficiency in control (CTL) and FXS NPCs at the G1 phase (PD-0332991 treatment) or at the G2/M border (Ro-3306 or nocodazole treatment). Cell cycle phase distribution after each treatment were analyzed by Fluorescence-Activated Cell Sorting (FACS) after DNA staining with propidium iodide (n = 4 cell lines/genotype; 2 replicates/cell line). On the left panel, representative DNA staining profiles after FACS analysis in each condition. **(C-D)** Protein levels of ERK and AKT activation following synchronization with **(C)** nocodazole (G2/M border) and **(D)** PD-0332991 (G1 phase) were analyzed by western-blot (top panel). On the bottom panel, quantification of phospho-ERK and phospho-AKT levels (n= 3 cell lines/genotype; 2 replicates/cell line). Results are indicated as mean ± SEM. ** p<0.01, ***p<0.0001, ****p<0.00001.

**Supplementary Figure 4**

**(A-B)** Quantification of neuron- and astrocyte-specific markers, respectively GAP43 **(A)** and GFAP **(B)** by quantitative PCR during the differentiation of CTL and FXS NPCs into neural culture (n = 4 cell lines/genotype, 2 replicates/cell line).

**Supplementary Figure 5**

**(A-B)** Western-blot of H3K79me2 global levels in NPCs following treatments with DOT1L inhibitors **(A)** within the same cell line and **(B)** across genotypes for each condition (n= 4 cell lines/genotype; 2 technical replicates/cell line) (left panels). On the right panels, quantification of proteins levels are indicated as mean ± SEM. *$p< 0.05$, ** $p<0.01$, ***$p<0.0001$, ****$p<0.00001$.

**Supplementary Figure 6**

**(A)** Schematic of the experimental design used to assay NPC differentiation. NPCs were pre-treated with 1 μM EPZ5676 or 1 μM SGC0946 for 72h, followed by differentiation in inhibitor-free conditions. **(B-C)** Quantification of the neuron-specific markers MAP2 **(B)** and GAP43 **(C)** by quantitative PCR during differentiation of CTL and FXS NPCs, pre-treated with DOT1L inhibitors, into neural cultures (n = 4 cell lines/genotype, 2 replicates/cell line) (left panel). The right panel shows the heatmap of p-values obtained from ANOVA analyses used to assess statistical significance between conditions. Data are presented as −log10(p-value). Values exceeding the significance threshold of 1.30 (corresponding to $p < 0.05$) are highlighted using a blue color gradient. **(D-E)** Heatmaps of p-values obtained from ANOVA analyses used to assess statistical significance between conditions for quantification of PLZF **(D)** and NeuN **(E)** expression levels. Data are presented as −log10(p-value). Values exceeding the significance threshold of 1.30 (corresponding to $p < 0.05$) are highlighted using a blue color gradient.

**Supplementary Figure 7**

**(A-B)** Heatmaps of p-values obtained from ANOVA analyses used to assess statistical significance between conditions for quantification of FABP7 **(A)** and ALDH1L1 **(B)** expression levels. Data are presented as −log10(p-value). Values exceeding the significance threshold of 1.30 (corresponding to $p < 0.05$) are highlighted using a blue color gradient. **(C)** Quantification of the astrocyte-specific marker GFAP by quantitative PCR during differentiation of CTL and FXS NPCs, pre-treated with DOT1L inhibitors, into neural cultures ($n = 4$ cell lines/genotype, 2 replicates/cell line) (left panel). The right panel shows the heatmap of p-values obtained from ANOVA analyses used to assess statistical significance between conditions. Data are presented as $-\log10$(p-value). Values exceeding the significance threshold of 1.30 (corresponding to $p < 0.05$) are highlighted using a blue color gradient.

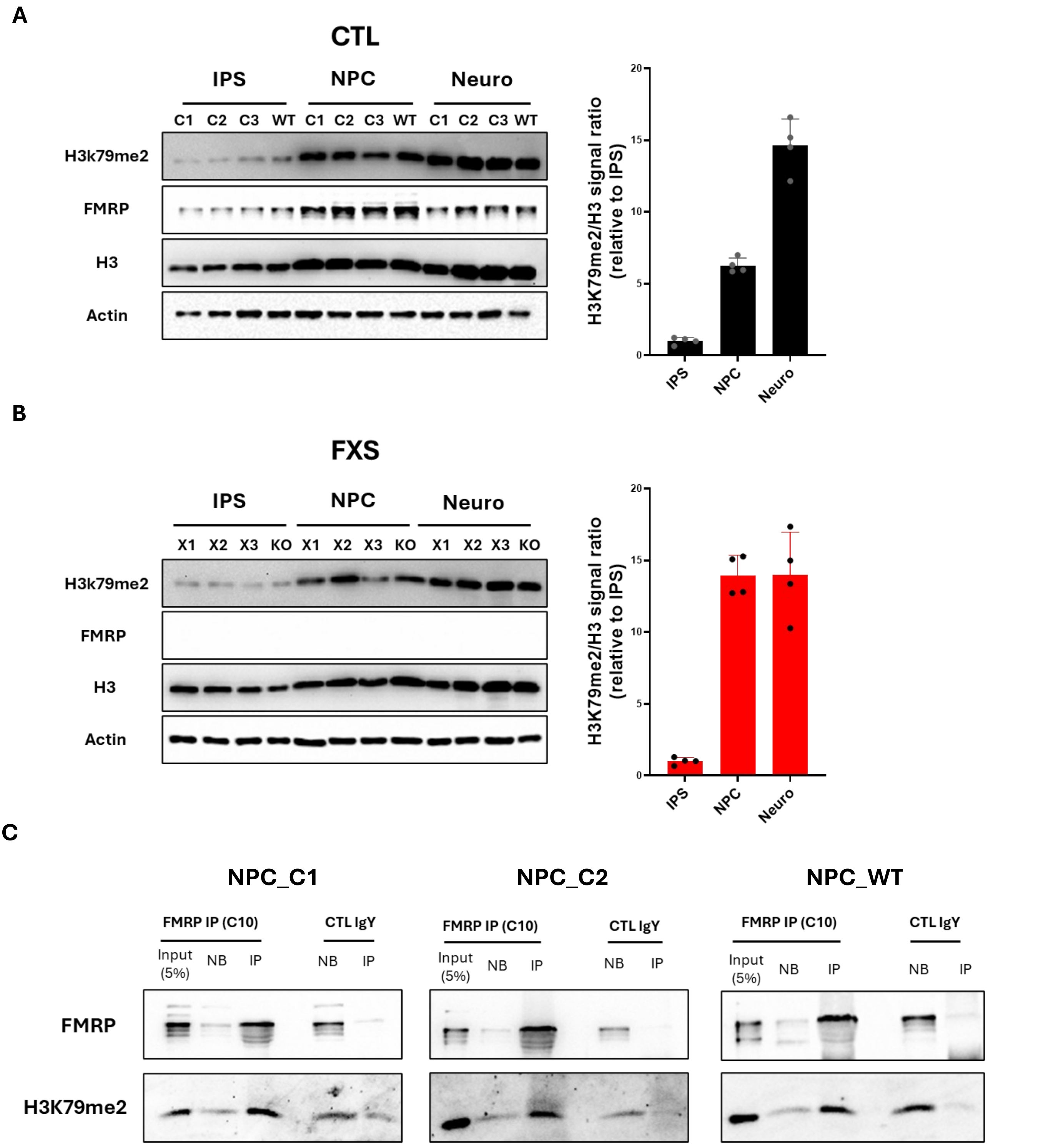


**Supplementary Figure 1**

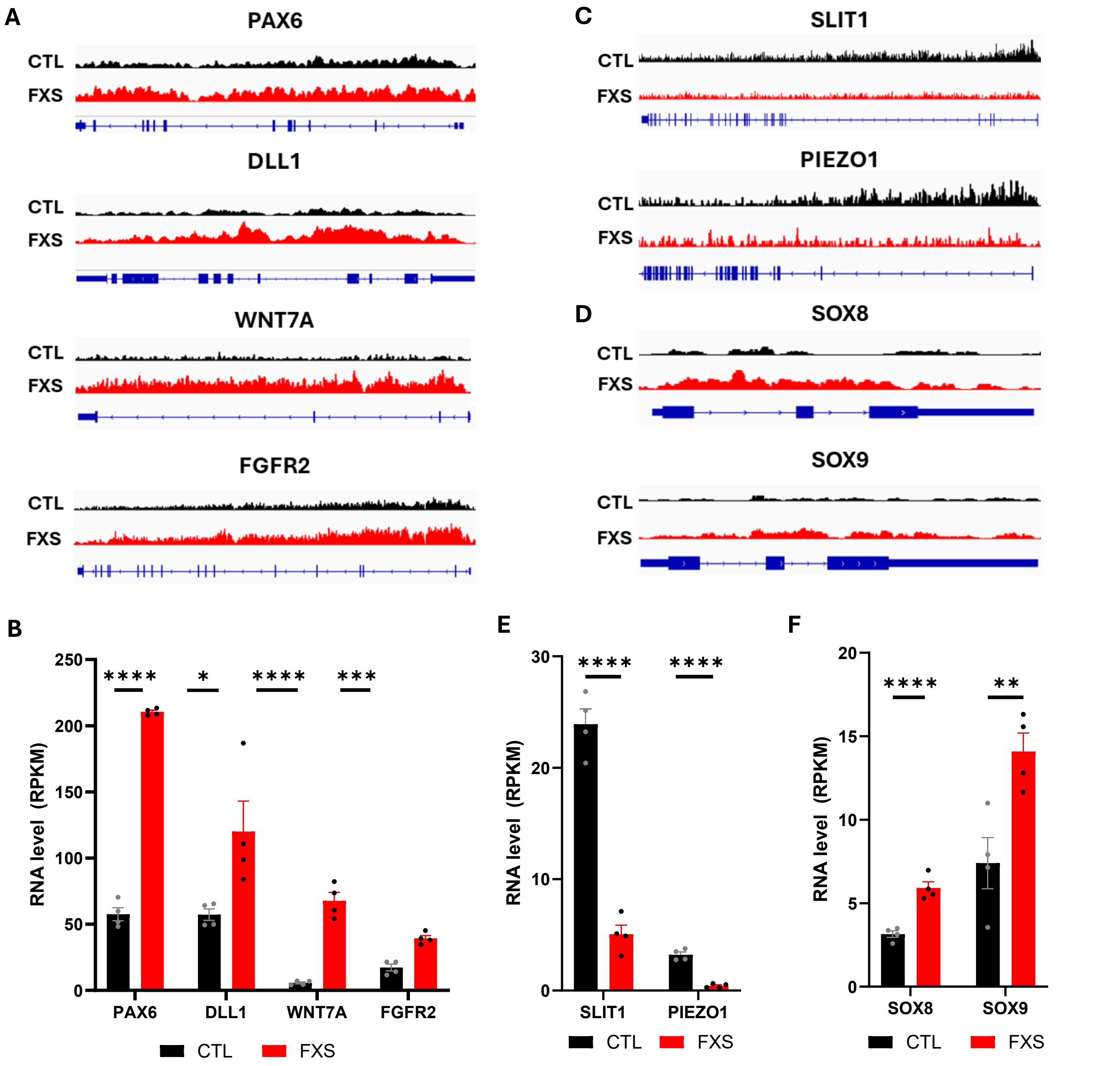


**Supplementary Figure 2**

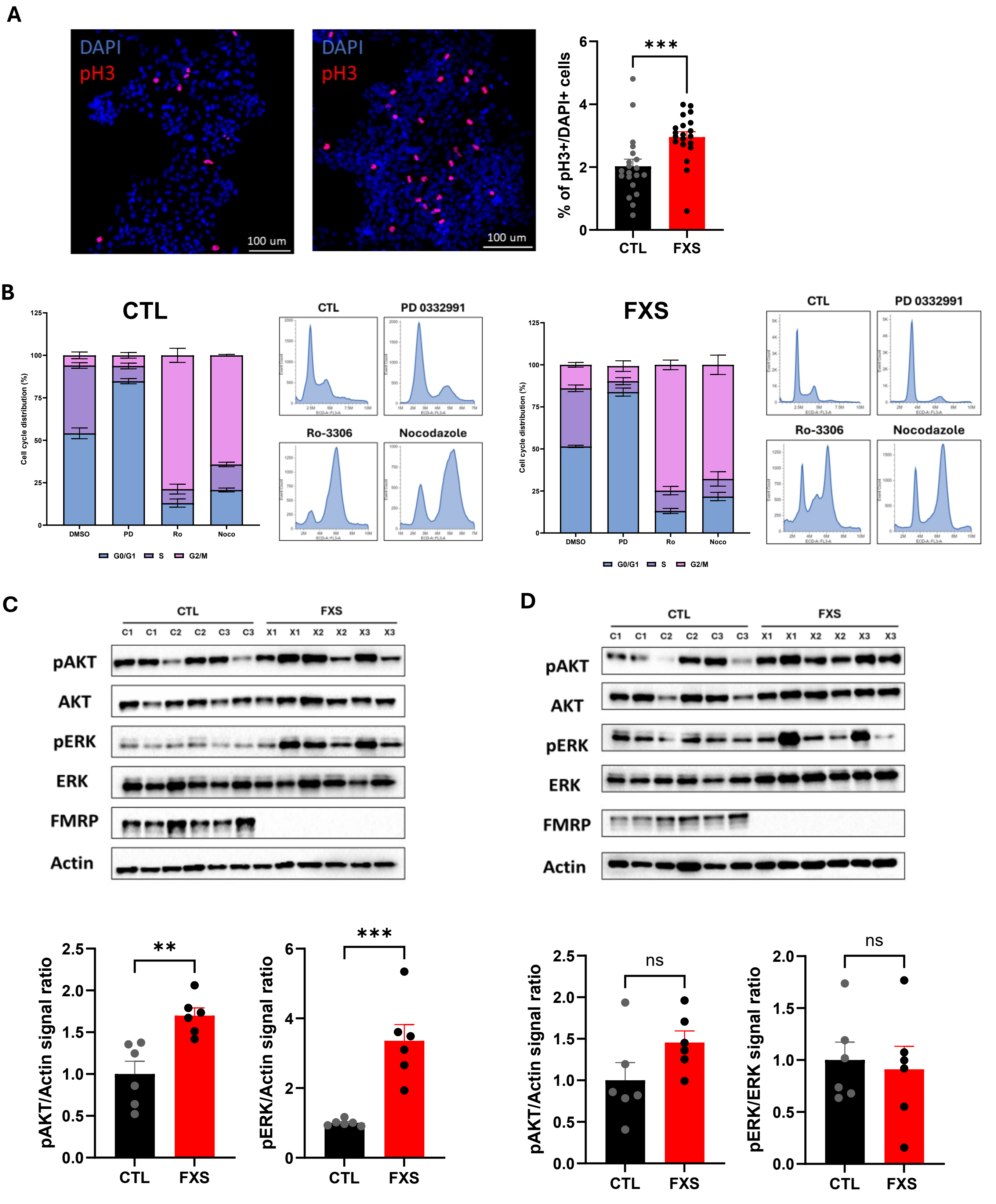


Supplementary Figure 3

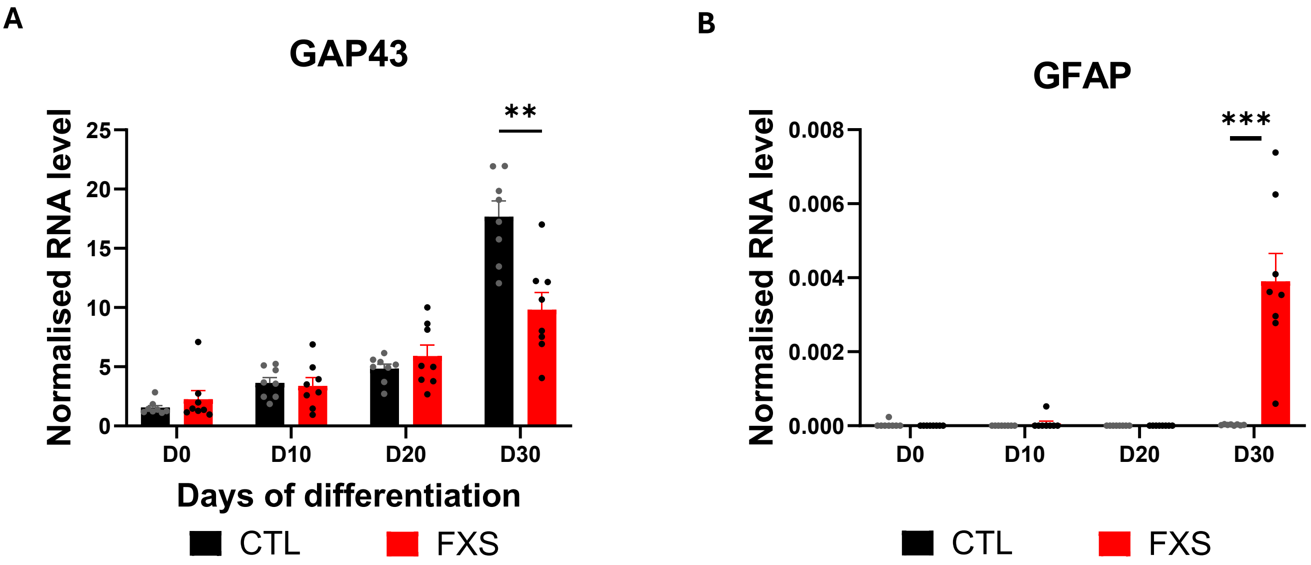


**Supplementary Figure 4**

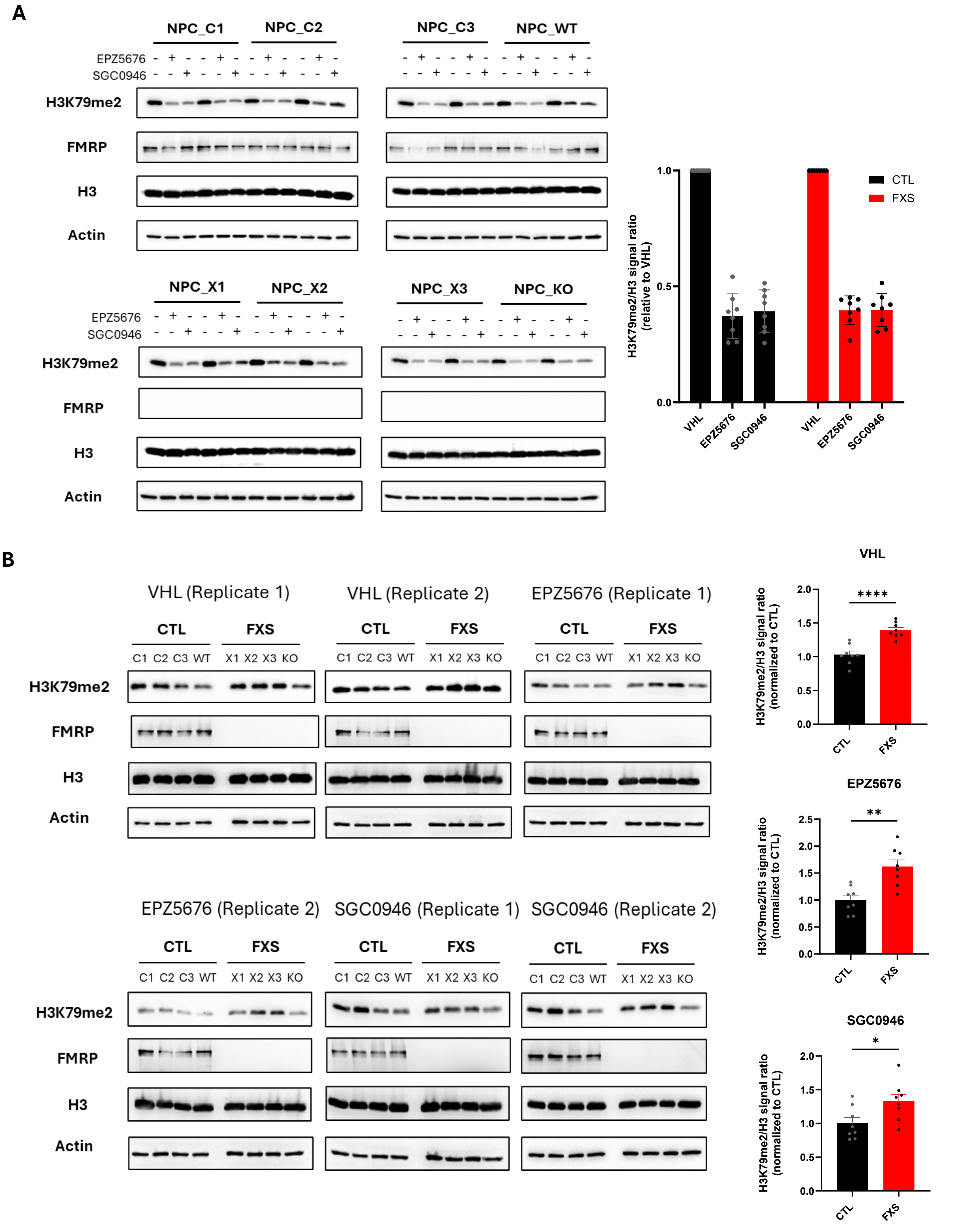


Supplementary Figure 5

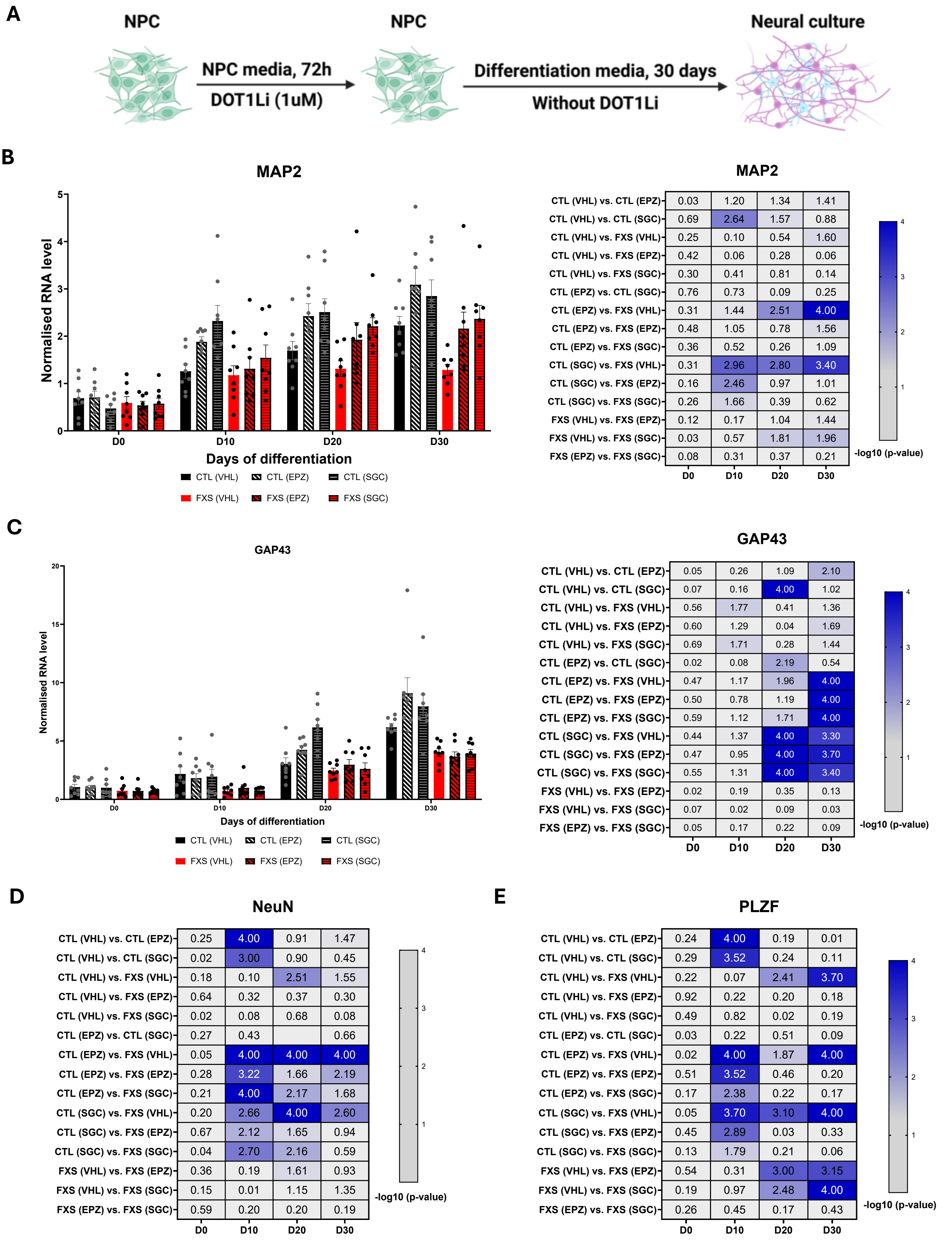


**Supplementary Figure 6**

**A**

#### FABP7

| Comparison | D0 | D10 | D20 | D30 |
|---|---|---|---|---|
| CTL (VHL) vs. CTL (EPZ) | 0.22 | 0.34 | 0.69 | 0.10 |
| CTL (VHL) vs. CTL (SGC) | 0.03 | 0.44 | 0.39 | 3.58e-003 |
| CTL (VHL) vs. FXS (VHL) | 0.18 | 0.06 | 0.44 | 2.33 |
| CTL (VHL) vs. FXS (EPZ) | 0.11 | 0.33 | 0.65 | 0.04 |
| CTL (VHL) vs. FXS (SGC) | 0.23 | 0.27 | 0.63 | 0.22 |
| CTL (EPZ) vs. CTL (SGC) | 0.27 | 1.00 | 0.19 | 0.10 |
| CTL (EPZ) vs. FXS (VHL) | 0.47 | 0.25 | 0.15 | 2.64 |
| CTL (EPZ) vs. FXS (EPZ) | 0.37 | 0.01 | 0.02 | 0.06 |
| CTL (EPZ) vs. FXS (SGC) | 0.54 | 0.05 | 0.03 | 0.36 |
| CTL (SGC) vs. FXS (VHL) | 0.14 | 0.55 | 0.03 | 2.31 |
| CTL (SGC) vs. FXS (EPZ) | 0.07 | 0.98 | 0.16 | 0.04 |
| CTL (SGC) vs. FXS (SGC) | 0.18 | 0.89 | 0.15 | 0.21 |
| FXS (VHL) vs. FXS (EPZ) | 0.06 | 0.24 | 0.12 | 2.46 |
| FXS (VHL) vs. FXS (SGC) | 0.03 | 0.19 | 0.11 | 1.75 |
| FXS (EPZ) vs. FXS (SGC) | 0.10 | 0.04 | 0.01 | 0.27 |

-log10 (p-value)

**B**

#### ALDH1L1

| Comparison | D0 | D10 | D20 | D30 |
|---|---|---|---|---|
| CTL (VHL) vs. CTL (EPZ) | 1.24 | 0.23 | 4.63e-003 | 0.43 |
| CTL (VHL) vs. CTL (SGC) | 1.09 | 0.10 | 0.08 | 0.80 |
| CTL (VHL) vs. FXS (VHL) | 0.14 | 0.38 | 4.85e-003 | 4.00 |
| CTL (VHL) vs. FXS (EPZ) | 1.36 | 0.13 | 0.06 | 1.54 |
| CTL (VHL) vs. FXS (SGC) | 1.50 | 0.33 | 0.07 | 1.87 |
| CTL (EPZ) vs. CTL (SGC) | 0.06 | 0.11 | 0.09 | 0.22 |
| CTL (EPZ) vs. FXS (VHL) | 0.93 | 0.10 | 0.01 | 4.00 |
| CTL (EPZ) vs. FXS (EPZ) | 0.05 | 0.08 | 0.07 | 0.73 |
| CTL (EPZ) vs. FXS (SGC) | 0.11 | 0.07 | 0.07 | 0.99 |
| CTL (SGC) vs. FXS (VHL) | 0.80 | 0.24 | 0.08 | 4.00 |
| CTL (SGC) vs. FXS (EPZ) | 0.11 | 0.03 | 0.02 | 0.38 |
| CTL (SGC) vs. FXS (SGC) | 0.18 | 0.19 | 0.01 | 0.58 |
| FXS (VHL) vs. FXS (EPZ) | 1.04 | 0.20 | 0.06 | 3.10 |
| FXS (VHL) vs. FXS (SGC) | 1.17 | 0.03 | 0.06 | 2.70 |
| FXS (EPZ) vs. FXS (SGC) | 0.05 | 0.16 | 0.01 | 0.12 |

-log10 (p-value)

**C**

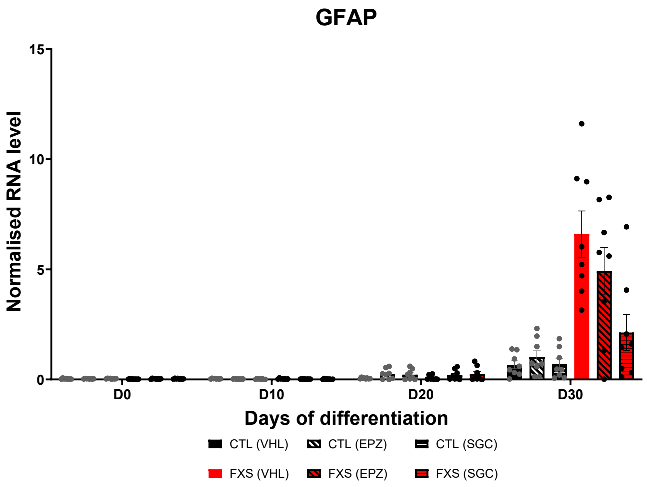


#### GFAP

| Comparison | D0 | D10 | D20 | D30 |
|---|---|---|---|---|
| CTL (VHL) vs. CTL (EPZ) | 0.13 | 0.85 | 1.08 | 0.12 |
| CTL (VHL) vs. CTL (SGC) | 0.48 | 1.42 | 0.89 | 0.01 |
| CTL (VHL) vs. FXS (VHL) | 0.84 | 0.42 | 0.07 | 4.00 |
| CTL (VHL) vs. FXS (EPZ) | 0.14 | 1.31 | 0.73 | 3.40 |
| CTL (VHL) vs. FXS (SGC) | 2.31e-003 | 1.56 | 1.03 | 3.40 |
| CTL (EPZ) vs. CTL (SGC) | 0.29 | 0.28 | 0.09 | 0.11 |
| CTL (EPZ) vs. FXS (VHL) | 1.12 | 0.27 | 0.91 | 4.00 |
| CTL (EPZ) vs. FXS (EPZ) | 0.30 | 0.22 | 0.18 | 3.05 |
| CTL (EPZ) vs. FXS (SGC) | 0.12 | 0.35 | 0.02 | 3.05 |
| CTL (SGC) vs. FXS (VHL) | 1.77 | 0.67 | 0.73 | 4.00 |
| CTL (SGC) vs. FXS (EPZ) | 0.72 | 0.04 | 0.08 | 3.40 |
| CTL (SGC) vs. FXS (SGC) | 0.48 | 0.05 | 0.06 | 3.40 |
| FXS (VHL) vs. FXS (EPZ) | 0.59 | 0.59 | 0.58 | 0.88 |
| FXS (VHL) vs. FXS (SGC) | 0.85 | 0.77 | 0.86 | 0.88 |
| FXS (EPZ) vs. FXS (SGC) | 0.14 | 0.09 | 0.15 | 4.34e-005 |

-log10 (p-value)

**Supplementary Figure 7**
